# TERRA R-loops trigger a switch in telomere maintenance towards break-induced replication and PrimPol-dependent repair

**DOI:** 10.1101/2025.01.09.632133

**Authors:** Suna In, Patricia Renck Nunes, Rita Valador Fernandes, Joachim Lingner

## Abstract

TERRA long noncoding RNAs associate with telomeres post transcription through base-pairing with telomeric DNA forming R-loop structures. TERRA regulates telomere maintenance but its exact modes of action remain unknown. Here, we induce TERRA transcription and R-loop formation in telomerase-expressing cells and determine that TERRA R-loop formation requires non-redundant functions of the RAD51 DNA recombinase and its enhancer RAD51AP1. TERRA R-loops interfere with semiconservative DNA replication promoting telomere maintenance by a homology-directed repair (HDR) mechanism known as Break Induced Replication (BIR), which ensures telomere maintenance in ALT cancer cells. In addition, TERRA induced PrimPol-dependent repair, which can initiate DNA synthesis de novo downstream of replication obstacles. PrimPol acts in parallel to BIR for telomere maintenance of TERRA over-expressing cells promoting their survival. Similarly, we find that PrimPol depletion is synthetic lethal with BIR deficiency in U2OS ALT cancer cells. Therefore, in the absence of other ALT-typical telomeric chromatin changes, TERRA R-loops per se are sufficient to induce ALT-typical telomere repair mechanisms.

## Introduction

Telomeric repeat containing RNAs (TERRA) are long noncoding RNAs that are transcribed from subtelomeric promoters towards chromosome ends (Azzalin *et al*, 2007; Feretzaki *et al*, 2019; Nergadze *et al*, 2009). TERRA molecules consist of chromosome end-specific subtelomeric sequences followed by telomeric UUAGGG repeats (Porro *et al*, 2014). TERRA can associate with chromosome ends through formation of RNA:DNA hybrid structures giving rise to R-loops, in which the telomeric G-rich DNA strand is displaced (Feretzaki *et al*, 2020). Intriguingly, TERRA R-loops can form post transcription in dependency of the RAD51 DNA recombinase (Feretzaki *et al*, 2020). RAD51 is bound to TERRA in nuclear extracts and it can catalyze R-loop formation in vitro suggesting that a RAD51-mediated homology search mechanism enables TERRA association with chromosome ends post transcription.

In telomerase-positive cancer cells and possibly most healthy human cells, TERRA is repressed in S phase probably to prevent the interference of TERRA R-loops with semiconservative DNA replication (Porro *et al*, 2010). However, the repression of TERRA in S phase is lost in alternative lengthening of telomeres (ALT) cancer cells (Flynn *et al*, 2015) promoting use of a telomerase-independent mechanism to counteract the end replication problem (Arora et al, 2014). Thus, TERRA provides oncogenic functions in ALT cancer (Kyriacou & Lingner, 2024). Increased TERRA expression and telomeric R-loops during S phase also occur at short telomeres in *Saccharomyces cerevisiae* cells from which telomerase was deleted (Graf *et al*, 2017). Significantly, activation of HDR at short telomeres required TERRA R-loops, which persisted in S phase slowing down telomere shortening. Similarly, TERRA R-loops were persistent in human cells that approached cellular senescence (Sze *et al*, 2023). In these cells, TERRA mediated sister telomere cohesion appeared to contribute either to telomere maintenance or telomere capping. Thus, in these examples, TERRA R-loops counteract short telomeres-induced cellular aging.

ALT is mostly seen in cancers of mesenchymal origin (Pickett & Reddel, 2015). ALT utilizes homology-directed repair (HDR) pathways to lengthen telomeres. Upon strand invasion of one telomere into another, unidirectional conservative telomeric DNA synthesis occurs through the engagement of a PCNA-DNA polymerase δ (Polδ) containing replisome (Dilley *et al*, 2016). This HDR pathway resembles break-induced replication (BIR), which has been thoroughly characterized in *S. cerevisiae* (Epum & Haber, 2022). However, HDR is repressed at intact healthy telomeres in normal cells and telomerase expressing cancer cells (Sfeir *et al*, 2010). To overcome HDR repression, ALT telomeres undergo extensive changes in telomeric DNA structure and protein composition. DNA damage response proteins and mediators of HDR appear and the histone variant H3.3 disappears at ALT telomeres (Déjardin & Kingston, 2009; Zhang *et al*, 2023; Kaminski *et al*, 2022). ALT telomeres also associate with PML proteins in subnuclear bodies referred to as ALT-associated PML bodies that mediate telomeric DNA synthesis. Elevated TERRA expression and increased TERRA R-loops are another feature of ALT cells and were proposed to promote recombination at ALT telomeres. Indeed, regulated levels of telomeric RNA-DNA hybrids at ALT telomeres are crucial to exert the ALT-typical DNA replication stress at telomeres and stimulate HDR (Arora *et al*, 2014; Silva *et al*, 2021, 2022, 2019). However, the extent to which the observed effects of TERRA are influenced by the unique composition of telomeric chromatin in ALT cells remained unclear.

In this study, we elucidate the roles of TERRA in telomere regulation by investigating the effects of increased TERRA transcription in non-ALT telomerase expressing cancer cells (HeLa). Using CRISPR activation to induce TERRA expression, we examine its impact on telomeric R-loop formation, DNA damage, and telomere fragility. We find that RAD51 and RAD51AP1 provide non-redundant roles to mediate TERRA R-loop formation. TERRA R-loops induce telomere damage, which is repaired either by break-induced replication or PrimPol-dependent repriming resolving replication stress and maintaining telomere integrity. We also find that PrimPol contributes to survival of U2OS ALT cancer cells. Thus, TERRA-induced telomere damage triggers DNA repair pathways that are required for cell survival of ALT cancer cells.

## Results

### TERRA overexpression leads to an increase in telomeric R-loops

To investigate the roles of TERRA in telomere regulation, we engineered HeLa cells for TERRA overexpression using CRISPR activation (Fig. 1A). To this end, we expressed four copies of VP16 (VP64), a strong transcription activator, in frame with catalytically dead Cas9 (dCas9) from a lentiviral vector. We also stably expressed either two guide RNAs which were designed to target TERRA promoter regions at multiple chromosome ends (Fig. 1A, Reagent and tools table), or a control guide RNA targeting the AAVS site (control cells) (Reagent and tools table). HeLa cells expressing TERRA promoter-targeting guide RNAs exhibited a 50-fold increase in TERRA expression compared to cells expressing the control guide RNA (Fig. 1B), an expression level which corresponds to a range that is typical for ALT cells. This overexpression was confirmed through RT-qPCR amplifying various subtelomeric TERRA sequences, demonstrating locus-specific TERRA induction (Fig. 1C). As expected, only subtelomeres with guide RNA binding sites showed an increase in TERRA levels (Fig. 1A,C). Consistent with VP16-mediated TERRA promoter activation, Chromatin Immunoprecipitation (ChIP) using an anti-acetyl-H4 antibody revealed increased H4 acetylation at guide RNA-targeted subtelomeres (Appendix Fig. S1A) but not within TTAGGG telomeric repeats (Appendix Fig. S1B).

**Figure 1.**
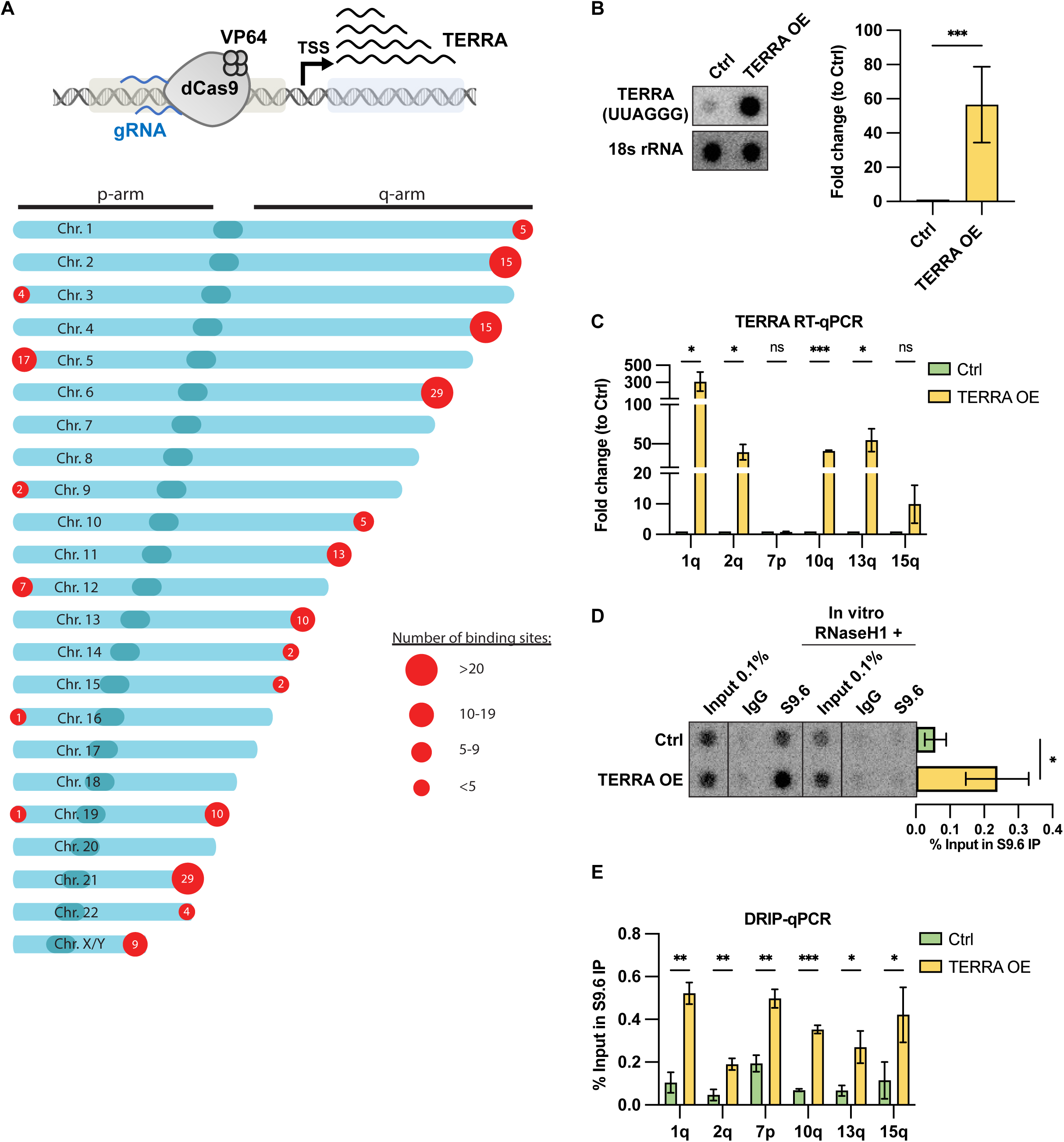
TERRA overexpression leads to an increase in telomeric R-loops. (A) Schematic representation of dCas9-VP64-induced expression of endogenous TERRA (top) and number of potential guide RNA binding sites near TERRA promoters (bottom). (B) RNA dot blot analysis in control and TERRA overexpressing HeLa cells containing an average telomere length of 10 kb (left) and quantification, plotted as fold change over Ctrl and normalized to 18s rRNA levels (right). Data represent mean ± s.d., from six independent biological replicates. Unpaired t-test was applied: *** P ≤ 0.001. (C) Quantification of TERRA levels by RT-qPCR analyzed using indicated subtelomeric primers, plotted as fold change over Ctrl and normalized to GAPDH RNA levels. Data represent mean ± s.d., from three independent biological replicates. Multiple unpaired t-test was applied: *** P ≤ 0.001, * P ≤ 0.05, ns indicates non-significance (P > 0.05). (D-E) DRIP assay using S9.6 antibody. DRIP samples and inputs were treated with RNase (DNase-free) and analyzed by DNA dot blot with a ^32^P-radiolabeled telomeric probe (D) or by qPCR with indicated subtelomeric primers (E). As a negative control, samples treated in vitro with RNaseH1 prior to immunoprecipitation were analyzed in parallel. (D) Data represent mean ± s.d., from three independent biological replicates. Unpaired t-test was applied: * P ≤ 0.05. (E) Data represent mean ± s.d., from three independent biological replicates. Multiple unpaired t-test was applied: *** P ≤ 0.001, ** P ≤ 0.01, * P ≤ 0.05.

Next, we performed DNA:RNA hybrid Immunoprecipitation (DRIP) using the S9.6 antibody (Aguilera & Ruzov, 2022) to examine telomeric R-loops upon TERRA induction. As a control for the specificity of the assay, we degraded the RNA moiety of R-loops in the isolated nucleic acids with RNaseH1 prior to immunoprecipitation, which abolished the DRIP signal (Fig. 1D). As expected, cells overexpressing TERRA showed higher levels of TERRA R-loops at telomeres compared to control cells (Fig. 1D,E). Notably, the 7p subtelomere, which does not overexpress TERRA, also formed more R-loops upon TERRA induction, supporting the notion that TERRA R-loops form post transcription in trans (Fig. 1E) (Feretzaki *et al*, 2020). Overall, the data demonstrate that the TERRA induction system in HeLa cells significantly increases endogenous TERRA expression and telomeric R-loops.

### RAD51 and RAD51AP1 are required for R-loop formation

Our recent study demonstrated that the DNA recombinase RAD51 promotes telomeric R-loop formation in HeLa cells (Feretzaki *et al*, 2020). However, in other works, the RAD51-interacting protein RAD51AP1 has been reported to promote TERRA R-loops in vitro (Yadav *et al*, 2022) and in ALT cells in order to facilitate the alternative lengthening of telomeres through its interaction with TERRA (Kaminski *et al*, 2022). We investigated whether RAD51AP1 plays a similar role in telomerase-positive cells. To this end, we depleted RAD51 and RAD51AP1 in HeLa cells (Fig. 2A) and performed DRIP analyses (Fig. 2B). Knockdown of either RAD51 or RAD51AP1 significantly suppressed telomeric R-loop formation in TERRA overexpressing cells (Fig. 2B). This indicates that both RAD51 and RAD51AP1 are crucial for the formation of telomeric R-loops under conditions of high TERRA expression. Notably, while the knockdown efficiency was similar in control cells, the depletion did not affect TERRA R-loop levels. It is possible that the majority of basal R-loops in the here-used HeLa cells with a relatively long telomere length of approximately 10 kb are formed co-transcriptionally, involving the invasion of nascent RNA into DNA in a RAD51-independent manner. Of note, HeLa cells with short telomeres contain more R-loops and show RAD51-dependent TERRA R-loop formation (Feretzaki *et al*, 2020). Taken together, our findings demonstrate that both RAD51 and RAD51AP1 are essential for telomeric R-loop formation in HeLa cells with elevated TERRA levels.

**Figure 2.**
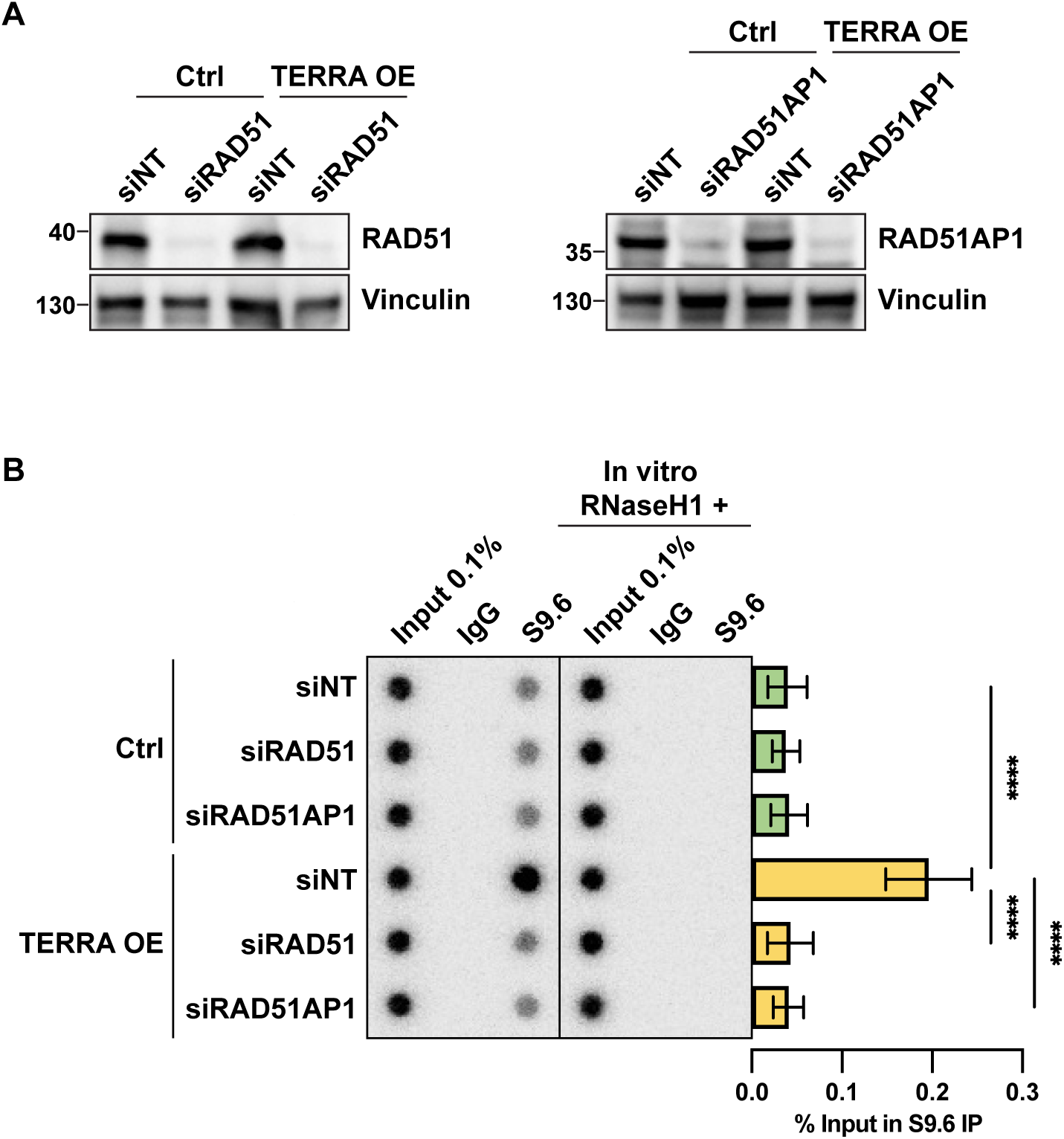
RAD51 and RAD51AP1 are required for TERRA R-loop formation. (A) Western blot analysis upon depletion of RAD51 (left) or RAD51AP1 (right) in control and TERRA overexpressing HeLa cells. (B) DRIP assay using S9.6 antibody. DRIP samples and inputs were treated with RNase (DNase-free) and analyzed by DNA dot blot with a ^32^P-radiolabeled telomeric probe. As a negative control, samples treated in vitro with RNaseH1 prior to immunoprecipitation were analyzed in parallel. Data represent mean ± s.d., from five independent biological replicates. Unpaired t-test was applied: * P ≤ 0.05. Data represent mean ± s.d., from three independent biological replicates. Multiple unpaired t-test was applied: **** P ≤ 0.0001.

### TERRA induces DNA damage at telomeres

R-loops are thought to potentially interfere with the replication machinery and they may contribute to DNA damage (García-Muse & Aguilera, 2019). To investigate this for TERRA R-loops, we assessed the frequency of telomere dysfunction-induced foci (TIFs) by performing immunofluorescence (IF) using an antibody against 53BP1, a marker of DNA damage, followed by telomeric fluorescent in-situ hybridization (FISH) (Fig. 3A,B). We evaluated the number of cells with more than five 53BP1 foci colocalizing with telomeric foci (Takai *et al*, 2003). Upon TERRA induction, there was a marked increase in the fraction of cells exhibiting more than five TIFs, comparable to the increase observed with zeocin treatment, which induces telomere damage as well as DNA double stranded breaks throughout the genome. Thus, TERRA overexpression alone induces significant DNA damage at telomeres (Fig. 3A,B). Additionally, cells with more than five non-telomeric 53BP1 foci were significantly increased by zeocin treatment, but not by TERRA overexpression (Appendix Fig. S2A,B). Thus, TERRA induces DNA damage specifically at telomeres. To determine whether TERRA-induced telomeric DNA damage results from R-loops, we ectopically expressed RNaseH1 (Appendix Fig. S2C) and assessed telomere TIFs. Notably, RNaseH1 overexpression completely rescued the increase in TIFs observed upon TERRA induction, while no effect was seen in control cells (Fig. 3C). These findings strongly suggest that TERRA R-loops are the primary drivers of DNA damage at telomeres.

**Figure 3.**
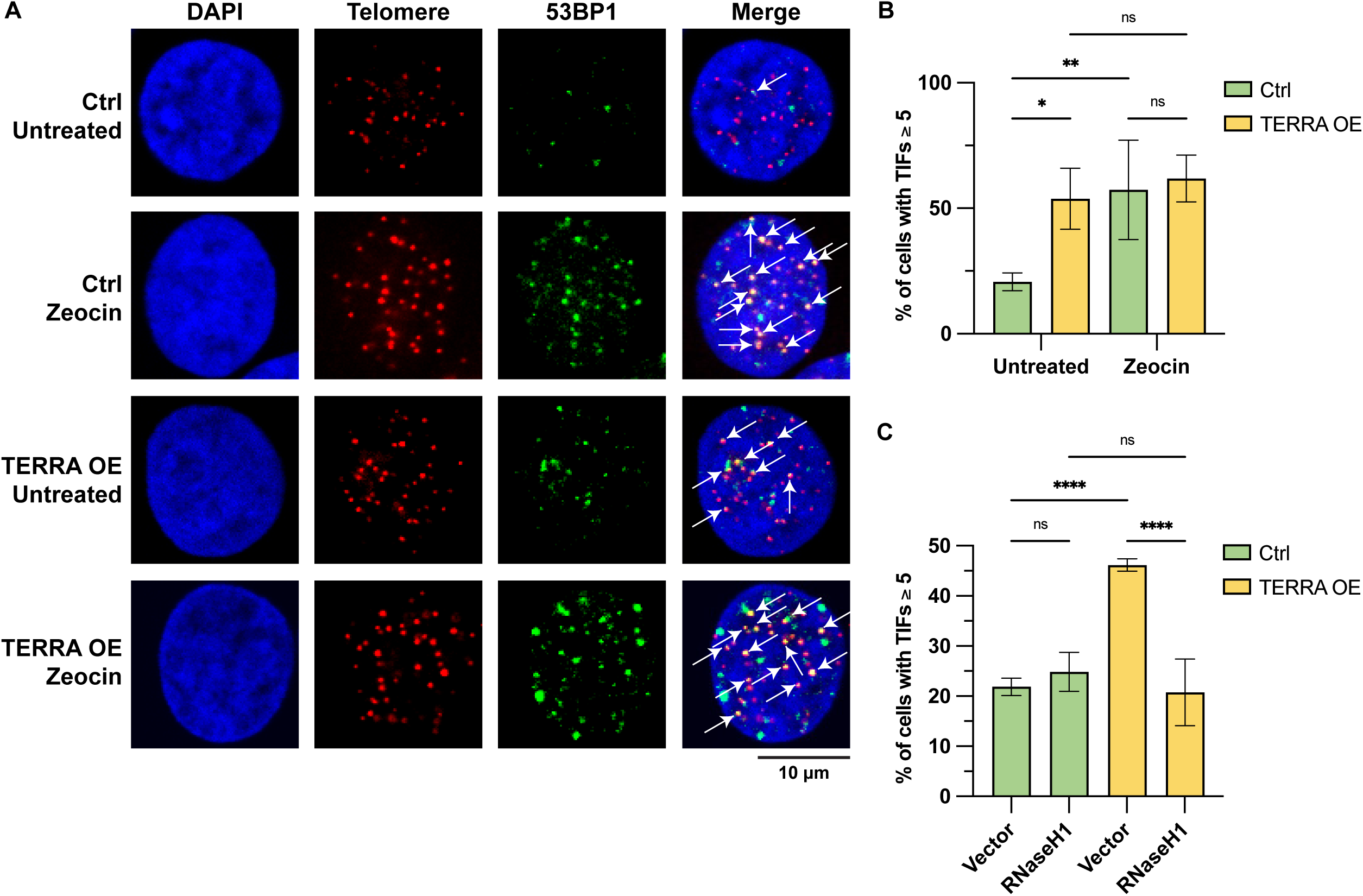
TERRA induces telomere dysfunction-induced foci (TIFs). (A) Representative images for detection of 53BP1 at telomeres in control and TERRA overexpressing HeLa cells. Immunofluorescence (IF) for 53BP1 (green) was combined with telomeric (CCCTAA)_3_-FISH (red) and DAPI staining (blue). (B) Quantification of the number of cells with ≥ 5 telomeres colocalizing with 53BP1. Data represent mean ± s.d., from three independent biological replicates. Two-way analysis of variance (ANOVA) with uncorrected Fisher’s least significant difference (LSD) test was applied: ** P ≤ 0.01, * P ≤ 0.05, ns indicates non-significance (P > 0.05). (C) Quantification of the number of cells with ≥ 5 telomeres colocalizing with 53BP1 upon RNaseH1 overexpression. Data represent mean ± s.d., from three independent biological replicates. Two-way analysis of variance (ANOVA) with uncorrected Fisher’s least significant difference (LSD) test was applied: **** P ≤ 0.0001, ns indicates non-significance (P > 0.05).

### TERRA R-loops directly induce telomere fragility

R-loops formed during S phase may potentially obstruct the replication machinery and contribute to DNA damage (García-Muse & Aguilera, 2019). Increased replication stress, induced by low doses of aphidicolin, which inhibits replicative DNA polymerases ⍺ and 𝛿, has been reported to display telomere fragility (Garcia-Exposito *et al*, 2016; Glousker & Lingner, 2021), characterized by smeared or doubled telomeric FISH signals on metaphase chromosomes (Fig. 4A). Several studies have reported an increase in telomere fragility when TERRA R-loop levels are elevated (Feretzaki *et al*, 2020; Fernandes & Lingner, 2023; Petti *et al*, 2019; Silva *et al*, 2019; Lin *et al*, 2021; Arora *et al*, 2014). However, these studies often indirectly affected TERRA R-loops by perturbing TERRA R-loop-regulating factors or were done in ALT cells. A study that most directly assessed TERRA R-loops on telomere fragility in HeLa cells utilized transgenic TERRA expressed from extrachromosomal plasmids (Feretzaki *et al*, 2020). Yet, to rule out the minor possibility that the PP7-coat protein fused to GFP (PCP-GFP) tag in transgenic TERRA influences the observed effects, it is crucial to evaluate telomere fragility driven specifically by endogenous TERRA R-loops.

**Figure 4.**
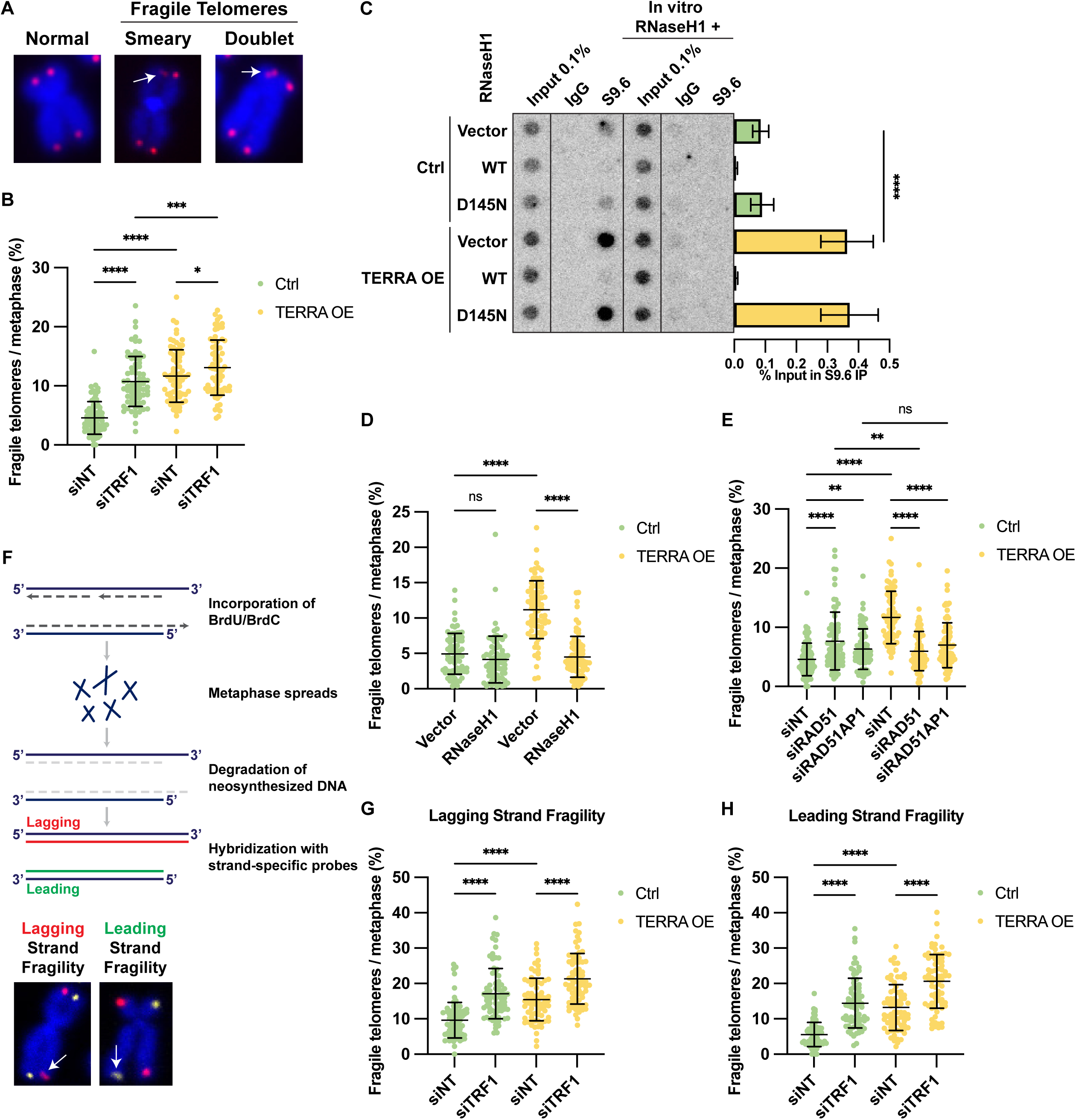
TERRA R-loops directly induce telomere fragility at both lagging and leading strands. (A) Representative images of metaphase spreads stained with telomeric (CCCTAA)_3_-FISH probe (red) and DAPI (blue). While normal telomeres show a single round signal at the end of each chromosome arm (left), fragile telomeres have smeary (middle) or multiple telomeric signals (right). White arrowheads indicate fragile telomeres. (B) Quantification of telomere fragility upon depletion of TRF1 in control and TERRA overexpressing HeLa cells. At least 25 metaphases were analyzed per condition per replicate, and three independent biological replicates were performed. Horizontal lines and error bars represent mean ± s.d. Two-way analysis of variance (ANOVA) with Tukey’s multiple comparisons test was applied: **** P ≤0.0001, *** P ≤ 0.001, * P ≤ 0.05. (C) DRIP (DNA:RNA hybrid immunoprecipitation) assay using S9.6 antibody. DRIP samples and inputs were treated with RNase (DNase-free) and analyzed by DNA dot blot with a ^32^P-radiolabeled telomeric probe. As a negative control, samples treated in vitro with RNaseH1 prior to immunoprecipitation were analyzed in parallel. Data represent mean ± s.d., from five independent biological replicates. Unpaired t-test was applied: * P ≤ 0.05 (D) Data represent mean ± s.d., from three independent biological replicates. Multiple unpaired t-test was applied: **** P ≤ 0.0001. (D) Quantification of telomere fragility upon overexpression of RNaseH1 in control and TERRA overexpressing HeLa cells. At least 25 metaphases were analyzed per condition per replicate, and three independent biological replicates were performed. Horizontal lines and error bars represent mean ± s.d. Two-way analysis of variance (ANOVA) with Tukey’s multiple comparisons test was applied: **** P ≤0.0001, ns indicates non-significance (P > 0.05). (E) Quantification of telomere fragility upon depletion of RAD51 or RAD51AP1 in control and TERRA overexpressing HeLa cells. At least 25 metaphases were analyzed per condition per replicate, and three independent biological replicates were performed. Horizontal lines and error bars represent mean ± s.d. Two-way analysis of variance (ANOVA) with Tukey’s multiple comparisons test was applied: **** P ≤0.0001, ** P ≤ 0.01, ns indicates non-significance (P > 0.05). (F) (Top) Schematic of the CO-FISH experiment. HeLa cells were incubated with BrdU and BrdC for 15 hours to ensure incorporation during a single round of replication. After metaphase enrichment via demecolcine treatment, cells were harvested, and metaphases were spread onto microscopic slides. Following RNaseA treatment and Hoechst staining, the newly synthesized DNA strands that incorporated BrdU and BrdC were degraded upon UV irradiation followed Exonuclease III digestion. The remaining parental DNA strands were then hybridized with strand-specific probes. The blue lines indicate parental strands, the dashed grey lines indicate BrdU/BrdC incorporated newly synthesized DNA strands, the red line indicates lagging strand-specific probes, and the green line indicates leading strand-specific probes. (Bottom) Representative images of metaphase spreads stained with TYE563-TeloC LNA probe (red), FAM-TeloG LNA probe (yellow), and DAPI (blue). White arrowheads indicate fragile telomeres. (G) Quantification of telomere fragility at lagging strand upon depletion of TRF1 in control and TERRA overexpressing HeLa cells with 30 kb telomeres. At least 25 metaphases were analyzed per condition per replicate, and three independent biological replicates were performed. Horizontal lines and error bars represent mean ± s.d. Two-way analysis of variance (ANOVA) with Tukey’s multiple comparisons test was applied: **** P ≤0.0001. (H) Quantification of telomere fragility at leading strand upon depletion of TRF1 in control and TERRA overexpressing HeLa cells with 30 kb telomeres. At least 25 metaphases were analyzed per condition per replicate, and three independent biological replicates were performed. Horizontal lines and error bars represent mean ± s.d. Two- way analysis of variance (ANOVA) with Tukey’s multiple comparisons test was applied: **** P ≤0.0001.

We analyzed telomere fragility upon overexpression of endogenous TERRA using the above introduced CRISPR-activation system and included TRF1 depletion (Appendix Fig. S3A), which is known to induce telomere fragility (Sfeir *et al*, 2009), as a positive control (Fig. 4B). The basal level of telomere fragility was observed to range between 3% and 5% (Fig. 4B). Similar to TRF1-depletion, TERRA overexpression led to an approximately 3-fold increase in telomere fragility to a frequency of 12% (Fig. 4B). The combination of TERRA overexpression and TRF1-depletion showed a partially additive effect (Fig. 4B). Thus, induction of endogenous TERRA in HeLa cells increases telomere fragility and TERRA exacerbates the telomere instability in TRF1 compromised cells.

To determine if the observed fragility was due to R-loops, we transiently overexpressed RNaseH1 (Appendix Fig. S3B), an enzyme that degrades the RNA moiety of R-loops (Fig. 4C). RNaseH1 overexpression suppressed the telomeric R-loops while a catalytically dead RNaseH1 (D145N) did not affect TERRA R-loops (Fig. 4C). At the same time, RNaseH1 overexpression completely suppressed the increased telomere fragility obtained upon TERRA overexpression (Fig. 4D). Of note, although DRIP analysis showed a further reduction of R-loops below basal levels upon RNaseH1 overexpression in control cells (Fig. 4C), telomere fragility did not drop below the basal level of roughly 3-5% (Fig. 4D). This suggests that the basal level of telomere fragility in HeLa cells stems mainly from other replication stress sources (G-quadruplex structures, unresolved t-loops, oxidative and other DNA damage, etc.). Indeed, the low endogenous TERRA levels in HeLa cells are further suppressed in S phase (Porro *et al*, 2010). As RNaseH1 depletion (Appendix Fig. S3C) in non-overexpressing cells increased telomere fragility to levels similar to TERRA overexpression (Appendix Fig. S3D), with no additional effect when TERRA was also induced (Appendix Fig. S3D), it seems that RNaseH1 suppresses R-loops in S phase cells. Overall, our experiments demonstrate that the increased telomere fragility seen in TERRA overexpressing cells is due to increased R-loop formation at telomeres and that RNaseH1 counteracts R-loops.

We also tested if RAD51 and RAD51AP1, both of which are required for TERRA R-loop formation, are required for telomere fragility. Strikingly, RAD51 or RAD51AP1 depletion suppressed the increase in fragility observed upon TERRA induction (Fig. 4E). In control cells, however, RAD51 and RAD51AP1 depletion caused a slight increase in fragility (Fig. 4E). This implies that RAD51 and RAD51AP1 may undertake several roles in regulating telomere fragility and maintenance; when TERRA levels are high, RAD51 and RAD51AP1 facilitate post-transcriptional TERRA R-loop formation, which then becomes the main source of telomere fragility. Conversely, when TERRA levels are low, depletion of these factors may perturb telomere replication and repair pathways that overcome the above-mentioned R-loop-independent replication obstacles. Overall, our results demonstrate that TERRA-induced telomere fragility requires RAD51 and RAD51AP1-dependent R-loops.

### TERRA overexpression increases telomere fragility at both leading and lagging strand telomeres

During telomere replication, the parental telomeric C-rich strand serves as template for leading strand synthesis while the G-strand serves as template for lagging strand synthesis. TERRA will base-pair with the C-rich strand when engaged in an R-loop structure. In these structures, the displaced single stranded G-rich telomeric DNA strand may form thermodynamically favored G-quadruplex structures (Yadav *et al*, 2022). Thus, in principle, the replication of both strands may be challenged by TERRA R-loops. Consistent with this notion, a recent study demonstrated that the accumulation of R-loops at telomeres induced by depletion of THOC subunits results in an increase of fragility of leading and lagging strand telomeres (Fernandes & Lingner, 2023). To assess the strand-specificity of telomere fragility upon TERRA induction, we performed telomeric chromosome-orientation FISH (CO-FISH) (Bailey *et al*, 1996). In brief, HeLa cells were incubated with BrdU and BrdC for 15 hours for incorporation of the modified nucleotides into the newly synthesized daughter strands (Fig. 4F). After spreading metaphase chromosomes on slides and staining with Hoechst dye, slides were UV irradiated, which preferentially damages the newly synthesized DNA strands containing the modified nucleotides. Following exonuclease III treatment, which degraded the nascent gap-containing DNA, the remaining G-rich parental strands were hybridized with 6-FAM-labeled LNA probes, while the parental C-rich strands were detected by hybridization with TYE563-labeled LNA probes (Fig. 4F). For optimal staining conditions, HeLa cells with 30 kb long telomeres were used to perform CO-FISH. Again, TRF1-depleted cells were used as a positive control. The basal level of telomere fragility detected by FISH was higher in the 30 kb telomere cells (Appendix Fig. S3E) than in cells with 10 kb telomeres (Fig. 4B), approximately 9%, as already reported (Fernandes & Lingner, 2023). Nonetheless, HeLa cells with 30 kb telomeres also showed increased telomere fragility upon TERRA induction or TRF1 depletion as seen in cells with 10 kb telomeres (Appendix Fig. S3E). Furthermore, CO-FISH experiments demonstrated that TERRA- induced telomere fragility occurred at both leading and lagging strand telomeres (Fig. 4G,H), aligning with findings from THOC subunit depletion studies (Fernandes & Lingner, 2023) as well as fragility obtained by TRF1-depletion (Fig. 4G,H) (Sfeir *et al*, 2009; Lee *et al*, 2018). Of note, TRF1 depletion also elevated the frequency of outsider telomeres (Majerska *et al*, 2018), in which the telomeric FISH signal is detached from chromosome ends (Appendix Fig. S3F), both at leading and lagging strands (Appendix Fig. S3F,G). The outsider phenotype was observed upon TRF1-depletion in HeLa cells (Fernandes & Lingner, 2023) but not in mouse cells from which the gene had been deleted (Sfeir *et al*, 2009; Yang *et al*, 2022). In contrast, TERRA overexpression did not increase the frequency of outsider telomeres, which therefore represents a distinct feature of human cells lacking TRF1 (Appendix Fig. S3G).

### BIR and PrimPol pathways act in parallel to repair TERRA R-loop mediated telomere damage

According to the prevalent view, telomere fragility is a phenomenon that occurs subsequent to replication stress. However, the precise mechanisms of its formation remain unclear (Glousker & Lingner, 2021). It is believed that the fragile appearance of telomeres stems from repair of stalled or collapsed replication forks in which single stranded DNA gaps are retained after repair and/or in which the repaired DNA is not fully condensed. TERRA R-loops may obstruct the replication machinery, leading to fork stalling and possibly fork collapse (García-Muse & Aguilera, 2019). One possible mechanism for restarting stalled forks might involve repriming downstream of the R-loop, a process that can be facilitated by PrimPol. The PrimPol polymerase has, unlike other DNA polymerases, the unique ability to initiate replication without the need of an RNA primer (Bianchi *et al*, 2013). BIR is another repair pathway which we suspected to repair TERRA R-loop dependent damage in HeLa cells, as it had already been linked to TERRA-induced replication stress and telomere maintenance in ALT cells . BIR is dependent on the POLD3 accessory subunit of DNA polymerase δ (Costantino *et al*, 2014). Furthermore, POLD3 depletion in mouse embryonic fibroblasts (MEFs) suppressed telomere fragility caused by knocking out the BLM RecQ helicase (Yang *et al*, 2020).

We tested whether single depletion of either PrimPol (Appendix Fig. S4A) or POLD3 (Appendix Fig. S4B) affected TERRA-induced telomere fragility (Fig. 5A,B). However, no significant changes of fragility were seen in TERRA overexpressing cells in which TERRA R-loops had become the major source of fragility (the increased fragility seen upon PrimPol or POLD3 in control cells may again, as discussed above for RAD51 and RAD51AP1, stem from roles of these factors in overcoming other replication stress sources). We therefore wondered whether the PrimPol and POLD3 pathways could compensate for each other and act in parallel pathways to overcome R-loop dependent telomere replication stress. Thus, when one pathway was inhibited, cells might use the other one to repair the telomere DNA damage. To test this hypothesis, we co-depleted PrimPol and POLD3. Strikingly, co-depletion of PrimPol with POLD3 resulted in a notable suppression of TERRA-induced telomere fragility in TERRA overexpressing cells (Fig. 5C). Of note, the reduction of fragility in TERRA overexpressing cells was not linked to effects on cell cycle distribution or R-loop frequency (Appendix Fig. S4C,D). From the decrease in telomere fragility upon co-depletion of PrimPol and POLD3 we conclude that both factors are involved in the repair of TERRA-induced telomere damage and the generation of telomere fragility.

**Figure 5.**
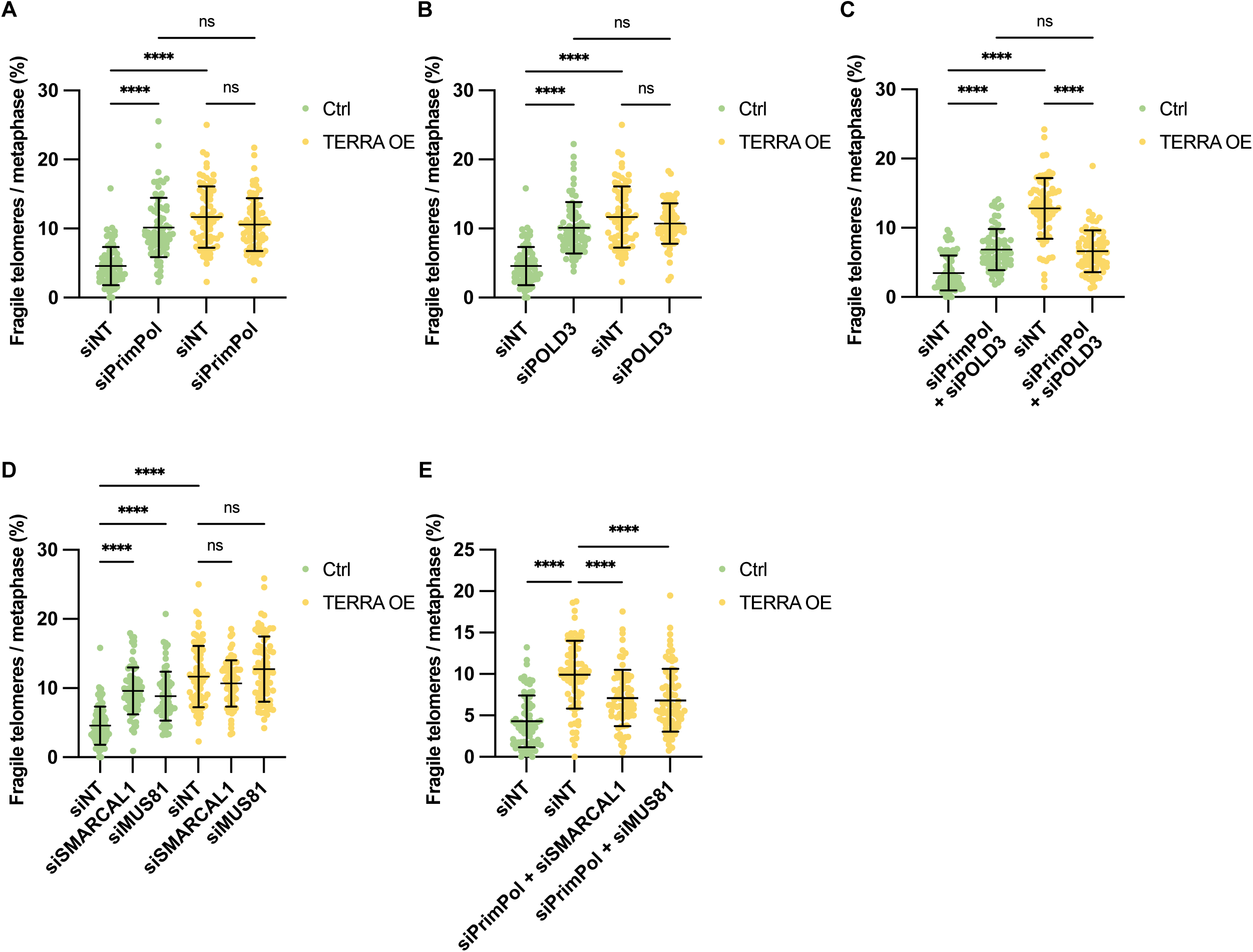
Repriming and BIR pathways act in parallel to repair TERRA R-loop mediated telomere damage. (A) Quantification of telomere fragility upon depletion of PrimPol in control and TERRA overexpressing HeLa cells. At least 25 metaphases were analyzed per condition per replicate, and three independent biological replicates were performed. Horizontal line and error bars represent mean ± s.d. Two-way analysis of variance (ANOVA) with Tukey’s multiple comparisons test was applied: **** P ≤0.0001, ns indicates non-significance (P > 0.05). (B) Quantification of telomere fragility upon depletion of POLD3 in control and TERRA overexpressing HeLa cells. At least 25 metaphases were analyzed per condition per replicate, and three independent biological replicates were performed. Horizontal lines and error bars represent mean ± s.d. Two-way analysis of variance (ANOVA) with Tukey’s multiple comparisons test was applied: **** P ≤0.0001, ns indicates non- significance (P > 0.05). (C) Quantification of telomere fragility upon co-depletion of PrimPol and POLD3 in control and TERRA overexpressing HeLa cells. At least 25 metaphases were analyzed per condition per replicate, and three independent biological replicates were performed. Horizontal lines and error bars represent mean ± s.d. Two-way analysis of variance (ANOVA) with Tukey’s multiple comparisons test was applied: **** P ≤0.0001, ns indicates non-significance (P > 0.05). (D) Quantification of telomere fragility upon depletion of PrimPol, SMARCAL1 or MUS81 in control and TERRA overexpressing HeLa cells. At least 25 metaphases were analyzed per condition per replicate, and three independent biological replicates were performed. Horizontal lines and error bars represent mean ± s.d. Two-way analysis of variance (ANOVA) with Tukey’s multiple comparisons test was applied: **** P ≤0.0001, ns indicates non-significance (P > 0.05). (E) Quantification of telomere fragility upon co-depletion of PrimPol and SMARCAL1 or MUS81 in control and TERRA overexpressing HeLa cells. At least 25 metaphases were analyzed per condition per replicate, and three independent biological replicates were performed. Horizontal lines and error bars represent mean ± s.d. Two-way analysis of variance (ANOVA) with Tukey’s multiple comparisons test was applied: **** P ≤0.0001.

To further characterize the involved pathways, we depleted the fork reversal enzyme SMARCAL1 (Appendix Fig. S4E) (Cox *et al*, 2016), depletion of which was reported to shift the balance toward PrimPol-dependent repriming (Quinet *et al*, 2020). We also depleted the fork cleavage enzyme MUS81 (Appendix Fig. S4F) (Sobinoff *et al*, 2017), which is required for replication restart after R-loop mediated fork stalling (Chappidi *et al*, 2020). Similar to POLD3 depletion, single depletion of SMARCAL1 or MUS81 did not affect TERRA-induced fragility (Fig. 5D). However, co-depletion of either SMARCAL1 or MUS81 with PrimPol significantly reduced TERRA-induced telomere fragility (Fig. 5E). As seen for PrimPol and POLD3 depletion, this reduction was not due to decreased R-loop levels (Appendix Fig. S4D). Taken together, our findings highlight the critical roles of PrimPol-dependent repriming and POLD3-dependent BIR in telomeric DNA repair following replication stress.

### Telomere repair by conservative DNA synthesis

In PrimPol mediated DNA repair, DNA synthesis should occur in a semiconservative manner using parental DNA as a template. In BIR, however, the broken telomeric DNA is elongated by conservative DNA synthesis. Upon strand invasion of the telomeric 3’ overhang, Polδ initiates repair DNA synthesis. The newly synthesized leading strand serves subsequently as a template for Okazaki strand synthesis (lagging strand) initiated by the polymerase alpha (Polα)-primase complex. In the CO-FISH protocol, the newly synthesized DNA strands are degraded (Fig. 4F). Thus, telomeres synthesized by conservative telomeric DNA synthesis should appear as truncated telomeres upon CO-FISH but not FISH analysis (Fig. 6A,B). We tested this possibility with HeLa cells containing an average telomere size of 30 kb. Consistent with the findings in HeLa cells with 10 kb telomeres, neither single depletion of PrimPol nor POLD3 significantly impacted TERRA-induced telomere fragility (Appendix Fig. S5). The FISH analysis also demonstrated that telomere loss events were very rare in TERRA overexpressing cells, independently of presence or absence of PrimPol and POLD3 (Fig. 6C). Significantly, however, in the CO-FISH analysis, telomere signal loss strongly increased upon TERRA overexpression, particularly at leading strand telomeres (Fig. 5C). In addition, telomere signal loss was augmented further upon PrimPol depletion, suggesting a shift towards BIR-mediated repair involving conservative DNA synthesis (Fig. 6C). POLD3 depletion showed a trend towards reduced telomere loss, which was, however, not statistically significant. Overall, we conclude that a significant portion of leading-strand telomeres, damaged due to the presence of TERRA R-loops, are repaired by conservative DNA synthesis. At lagging strand telomeres, the CO-FISH analysis also indicates conservative DNA replication-mediated repair upon TERRA overexpression, which was, however, occurring at lower levels (Fig. 6C). This suggests that the increased formation of G-quadruplex structures, prompted by TERRA R-loops, also necessitates the BIR-mediated conservative repair pathway (Yang *et al*, 2020).

**Figure 6.**
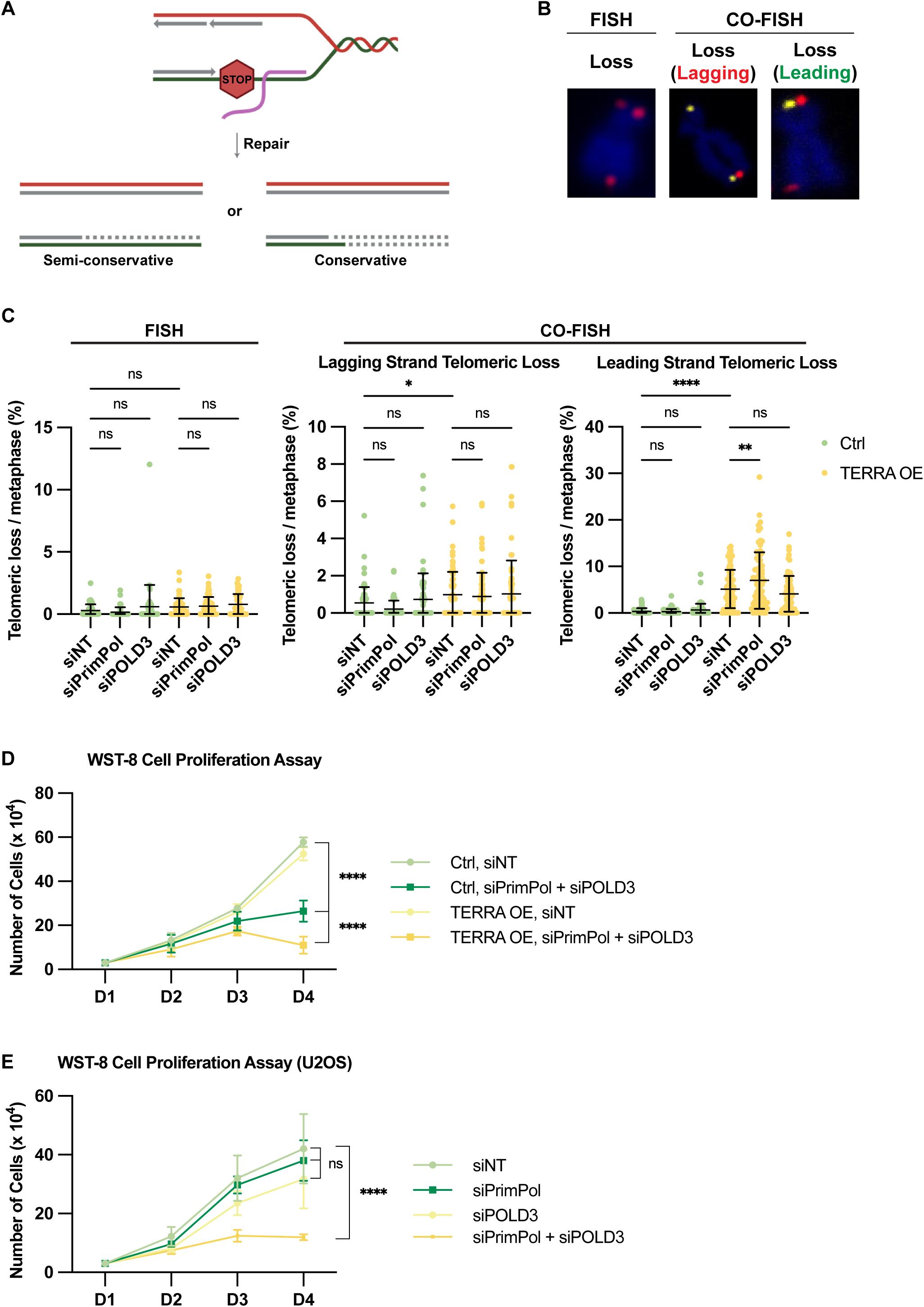
TERRA-induced DNA damage requires conservative repair pathway. (A) Schematic representation of two possible scenarios that occur during DNA repair caused by TERRA R-loops. The red line represents the parental strand for lagging strand synthesis, while the green line represents the parental strand for leading strand synthesis. Solid grey lines indicate newly synthesized DNA during replication, and dashed grey lines represent newly synthesized DNA during repair. Both the solid and dashed grey lines are degraded and therefore not detected in CO-FISH experiments. (B) Representative images of telomeric loss by FISH (left) or CO-FISH (middle and right). Metaphase spreads were stained with telomeric (CCCTAA)_3_-FISH probe (red) and DAPI (blue) (left), with TYE563-TeloC LNA probe (red), FAM-TeloG LNA probe (yellow), and DAPI (blue) (middle and right). (C) Quantification of telomeric loss upon depletion of PrimPol or POLD3 in control and TERRA overexpressing HeLa cells with 30 kb telomeres. Metaphases from the same samples were analyzed by FISH (left) and CO-FISH (middle and right). At least 25 metaphases were analyzed per condition per replicate, and three independent biological replicates were performed. Horizontal lines and error bars represent mean ± s.d. Two-way analysis of variance (ANOVA) with Tukey’s multiple comparisons test was applied: **** P ≤0.0001, ** P ≤ 0.01, * P ≤ 0.05, ns indicates non-significance (P > 0.05). (D) WST-8 cell proliferation assay upon co-depletion of PrimPol and POLD3 in control and TERRA overexpressing HeLa cells. After seeding, cell viability was measured every 24 hours. siRNA transfection was performed at 24 hr. Two-way analysis of variance (ANOVA) with Tukey’s multiple comparisons test was applied: **** P ≤0.0001. (E) WST-8 cell proliferation assay upon single or co-depletion of PrimPol and/or POLD3 in U2OS cells. After seeding, cell viability was measured every 24 hours. siRNA transfection was performed at 24 hr. Two-way analysis of variance (ANOVA) with Tukey’s multiple comparisons test was applied: **** P ≤0.0001, ns indicates non-significance (P > 0.05).

Finally, we tested the importance of PrimPol and POLD3 for the survival of TERRA overexpressing cells. Significantly, co-depletion of PrimPol and POLD3 impaired cell growth in TERRA overexpressing cells much more severely than in control cells (Fig. 6D). This indicates that both repair pathways make crucial contributions to the survival of cells suffering from TERRA R-loop-induced telomere damage. Strikingly, co-depletion of PrimPol and POLD3 significantly impaired cell growth of U2OS cells, whereas single depletion of either factor did not show a significant impact (Fig. 6E). This further supports the critical interplay between PrimPol and POLD3 in supporting survival mechanisms required in cells with high TERRA levels.

## Discussion

The consequences and functions of TERRA R-loops have been studied in previous work mostly in the context of short and damaged telomeres during cellular senescence or at ALT telomeres, which contain a dramatically altered telomeric chromatin composition. In this paper, we investigate the formation and consequences of TERRA R-loops at healthy telomeres with normal length by inducing expression of endogenous TERRA with CRISPR activation. The strong increase in telomere fragility upon TERRA induction allowed us to distinguish the mechanisms related to TERRA R-loops from other sources of damage. Thus, important biological effects and functions at telomeres could be directly assigned to TERRA. Our work provides a framework explaining TERRA R-loop biology, which consists of multiple steps (Fig. 7). First, elevated TERRA transcription leads to an increase in telomeric R-loops that not only form at the telomeres from which TERRA is expressed but also at telomeres in trans post transcription. Post-transcriptional TERRA R-loop formation requires both the RAD51 DNA recombinase and the RAD51 interacting protein RAD51AP1, which has been implicated in assisting HDR during synaptic complex formation and strand invasion (Pires *et al*, 2017). More recently, RAD51AP1 has also been demonstrated to be required for TERRA R-loop formation and telomere maintenance in ALT cancer cells (Barroso-González *et al*, 2019). Second, upon R-loop formation, TERRA R-loops interfere with the semiconservative DNA replication machinery leading to telomere damage and telomere fragility. Indeed, the telomere fragility was alleviated when TERRA R-loops did not form because of RAD51 or RAD51AP1 depletion or when they were destroyed by RNaseH1 overexpression. Third, the replication interference by TERRA R-loops leads to replication fork damage followed by DNA repair. Two pathways function in parallel to evade the TERRA-R-loop-induced replication blockade. PrimPol, which can reprime and therefore restart DNA synthesis downstream of obstacles, can overcome TERRA R-loop-mediated fork stalling. It remains uncertain if residual TERRA may serve as a primer for leading strand synthesis. Alternatively, POLD3-dependent repair can kick in to repair TERRA R-loop-generated damage. The POLD3 accessory subunit of DNA polymerase δ is essential for BIR.

**Figure 7.**
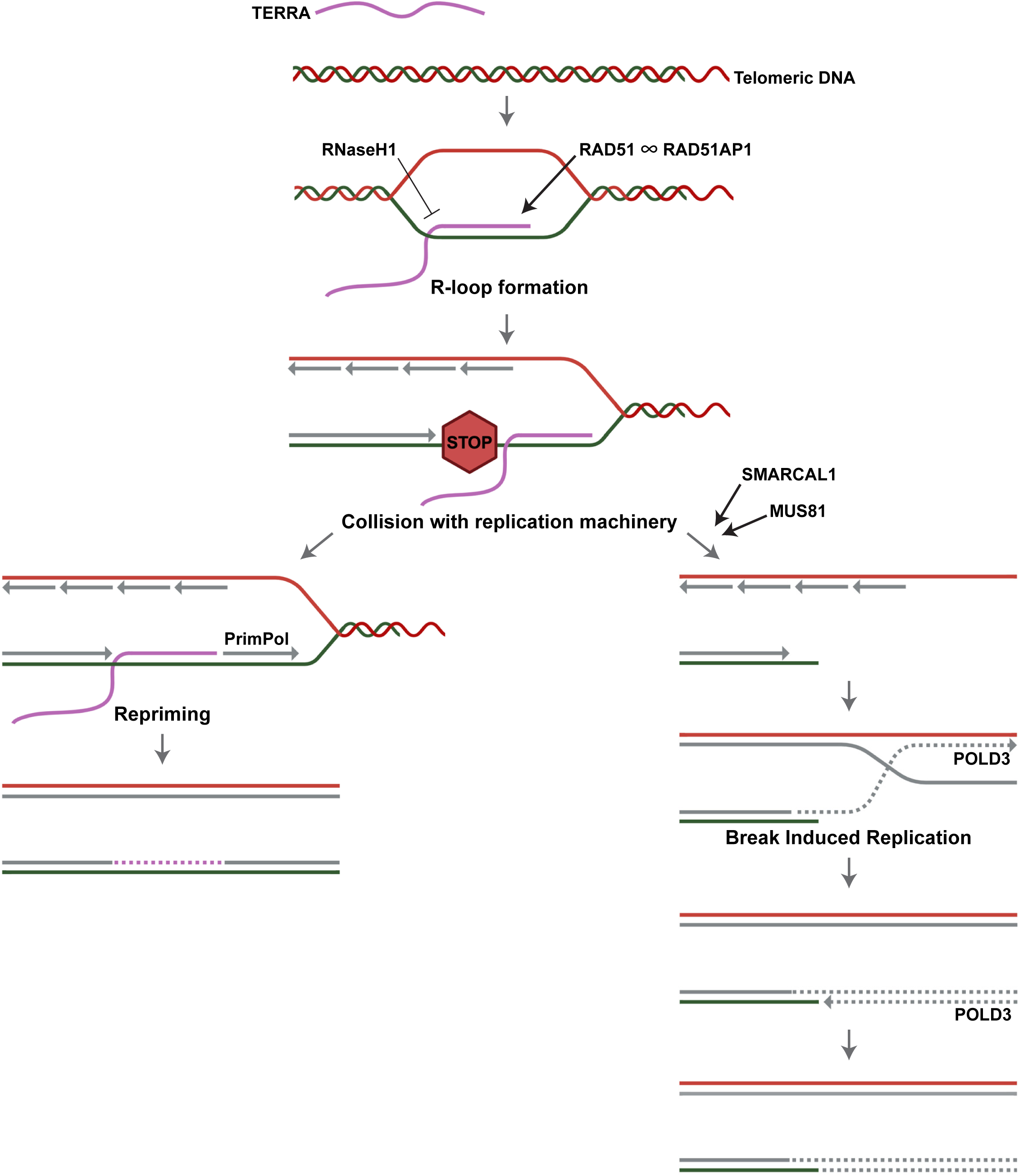
Model for TERRA R-loop formation, replication interference and repair. RAD51 and RAD51AP1 are both required for the formation of TERRA R-loops post transcription. TERRA R-loops can be destroyed by RNaseH1. In S phase, TERRA R-loops interfere with the DNA replication fork progression giving rise to telomere damage and repair. Two main pathways are involved in the repair of TERRA R-loop damaged DNA. Either, PrimPol mediates fork restart downstream of TERRA R-loops or the truncated telomeric DNA is repaired by POLD3-dependent BIR. While the specific roles of SMARCAL1 and MUS81 in these processes have not been precisely defined, they function in parallel to PrimPol and may prepare the replication-impaired telomeric DNA for strand invasion and repair synthesis. Remaining gaps due to incomplete DNA repair synthesis may give rise to telomere fragility. Solid grey lines represent newly synthesized DNA by semiconservative DNA replication. Purple dashed lines denote gaps which may remain from TERRA R-loops. Grey dashed lines represent newly synthesized DNA via BIR. This model does only depict repair pathways at leading strand telomeres. Elevated G-quadruplex formation at single-stranded G-rich strands, driven by TERRA R-loops, along with subsequent repair by BIR may also contribute to enhanced fragility at the lagging strand.

Furthermore, the SMARCAL1 ATP-dependent strand annealing helicase is involved (Cox *et al*, 2016). SMARCAL1 catalyzes fork remodeling and/or fork reversal and, as such, appears to mediate repair stemming from R-loop induced damage. The MUS81 nuclease is also required for the repair. MUS81 contributes to ALT-mediated DNA synthesis (Lu *et al*, 2024). MUS81 is part of a specialized endo-nucleolytic complex known as SMX (SLX1-4, MUS81-EME1, XPF-ERCC1) (Sobinoff *et al*, 2017) and may cleave impeded and rearranged recombination intermediates possibly to generate telomeric DNA substrates for strand invasion. The POLD3 dependent repair involves conservative DNA synthesis consistent with the BIR mechanism. The CO-FISH data indicate that BIR is mostly engaged at leading strand telomeres in TERRA overexpressing cells. The crucial importance of the POLD3 and PrimPol repair pathways for TERRA R-loop containing telomeres is apparent as concomitant impairment of POLD3 and PrimPol caused massive cell death in TERRA overexpressing cells.

Overall, our results demonstrate that TERRA R-loops are sufficient to induce replication stress at telomeres, which is followed by telomere maintenance by BIR. This mechanism may become particularly important at short telomeres, especially in telomerase-negative cells, to prevent premature senescence (Graf *et al*, 2017; Sze *et al*, 2023). Indeed, we did not observe any changes in telomere length following TERRA induction in HeLa cells with 10 kb telomeres after several weeks of growth (data not shown). In telomerase-negative ALT cancer, the exact genetic and epigenetic alterations that lead to the activation of ALT-telomere maintenance mechanisms are unknown. We searched in TERRA overexpressing HeLa cells for ALT-typical characteristics including enhanced sister chromatid exchange frequency, formation of C-circles and ALT-specific PML bodies. However, none of these features was induced indicating that TERRA has specific effects on telomere replication and repair (data not shown). Thus, other genetic or epigenetic changes are required to induce a complete ALT phenotype. It has been demonstrated that experimental depletion of histone chaperones ASF1a and ASF1b in human cells induced ALT features (O’Sullivan *et al*, 2014). ATRX-loss is also seen frequently in ALT cancer cells contributing to telomeric chromatin changes and TERRA expression (Heaphy *et al*, 2011). Our results indicate that increased TERRA R-loops are sufficient to induce BIR suggesting that their establishment may represent a key event in ALT cancer cell evolution to overcome cell death during replicative cell crisis.

Intriguingly, during cell crisis, TERRA is upregulated at TRF2-depleted critically short telomeres acting as signaling molecule in the cytoplasm and triggering innate immune response-dependent autophagic cell death (Nassour *et al*, 2023). Thus, upregulated TERRA promotes cell death during crisis acting as a tumor suppressor. TERRA upregulation during cell crisis, however, may become a two-edged sword and acquire oncogenic function when deviated into R-loops to initiate the BIR-mediated telomere maintenance mechanisms of ALT cells.

## Methods

### Reagents and tools table

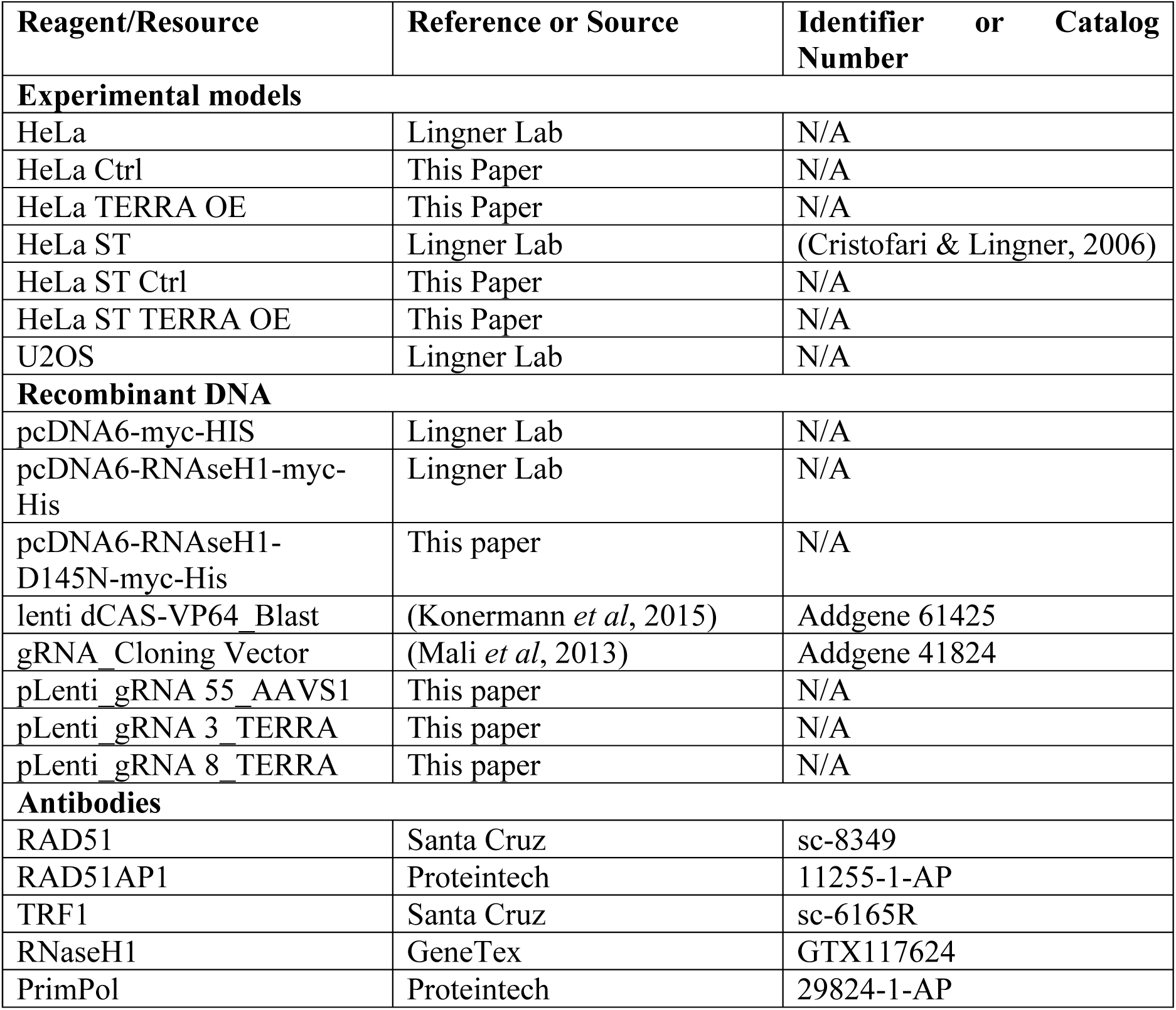
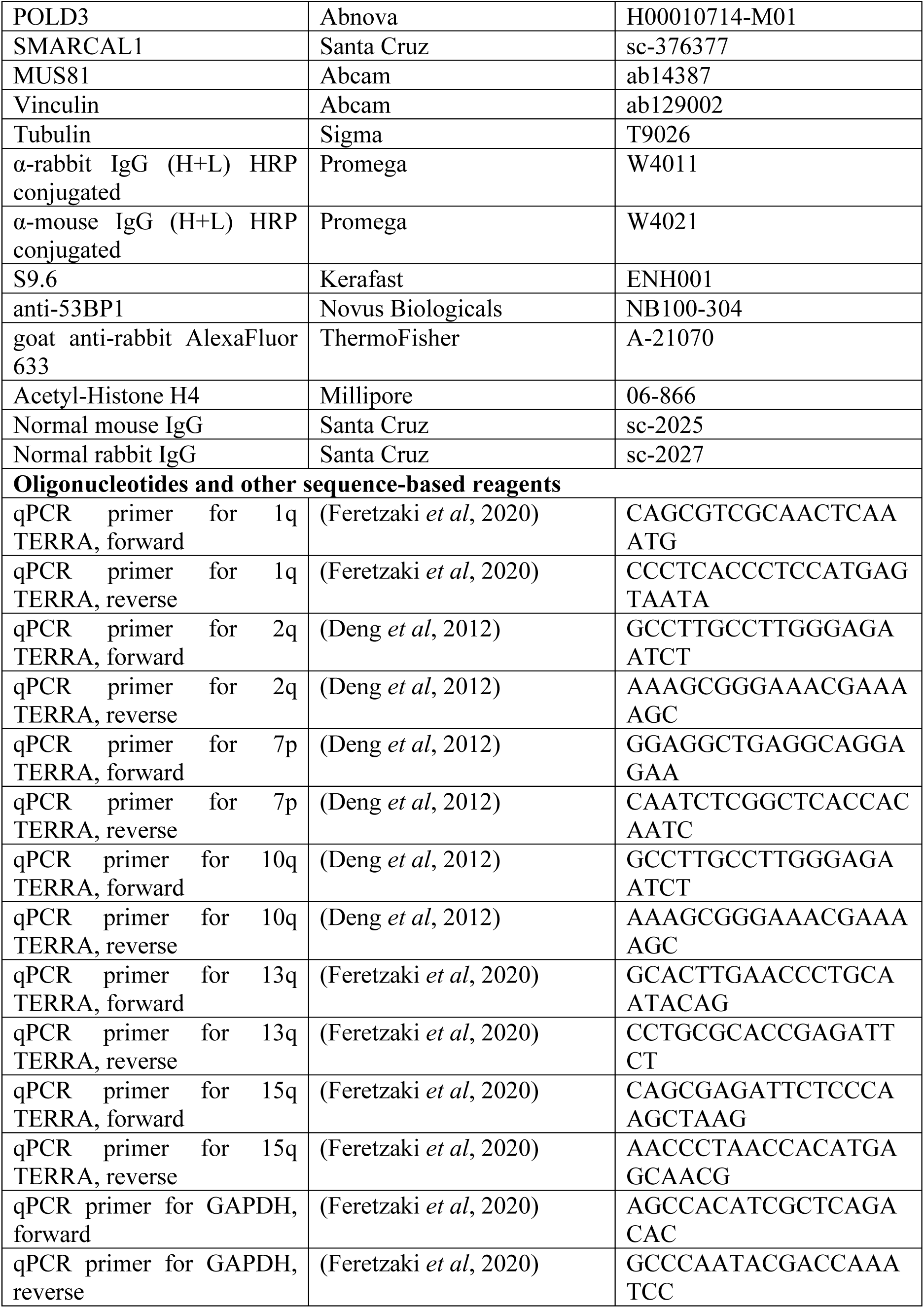
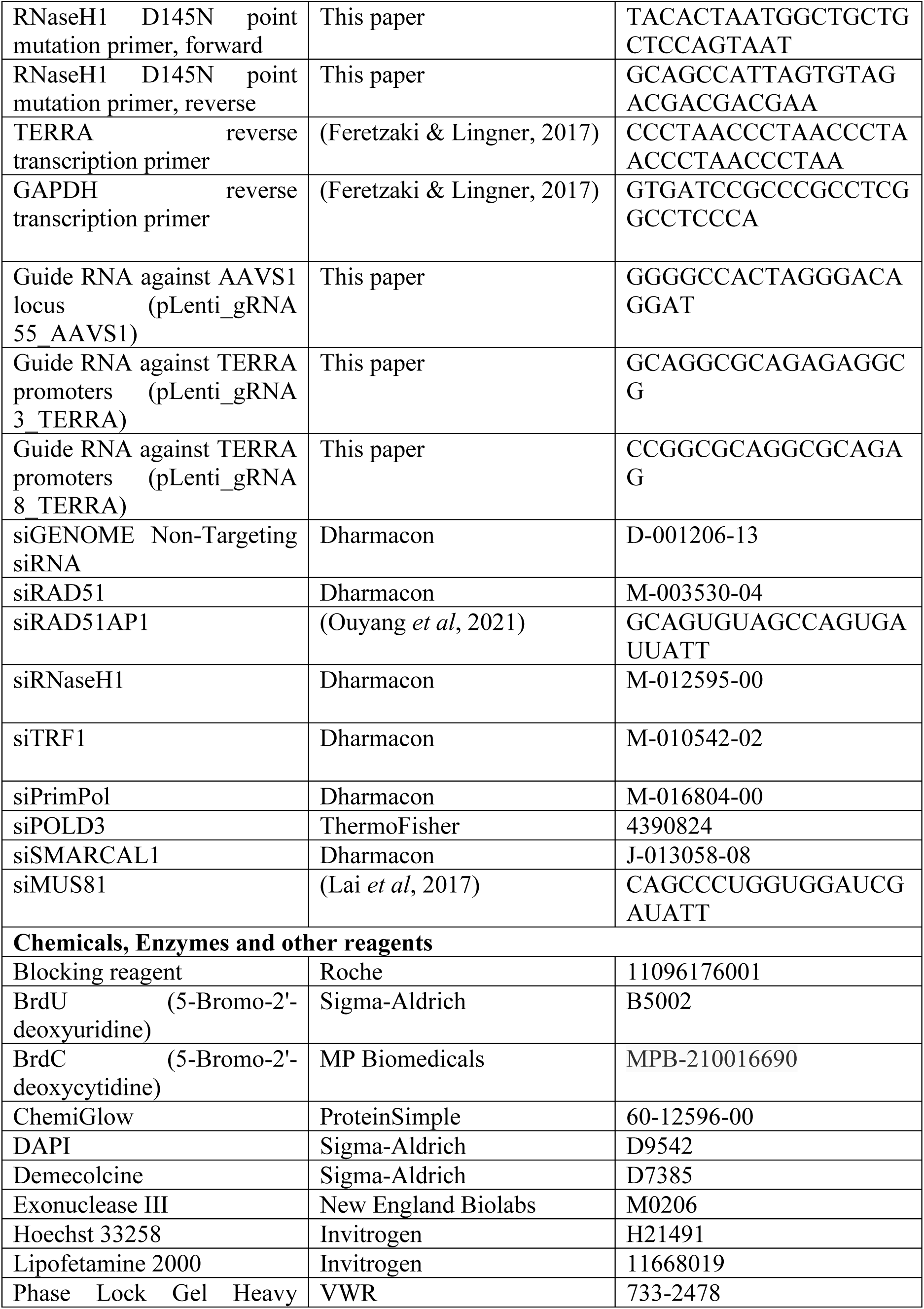
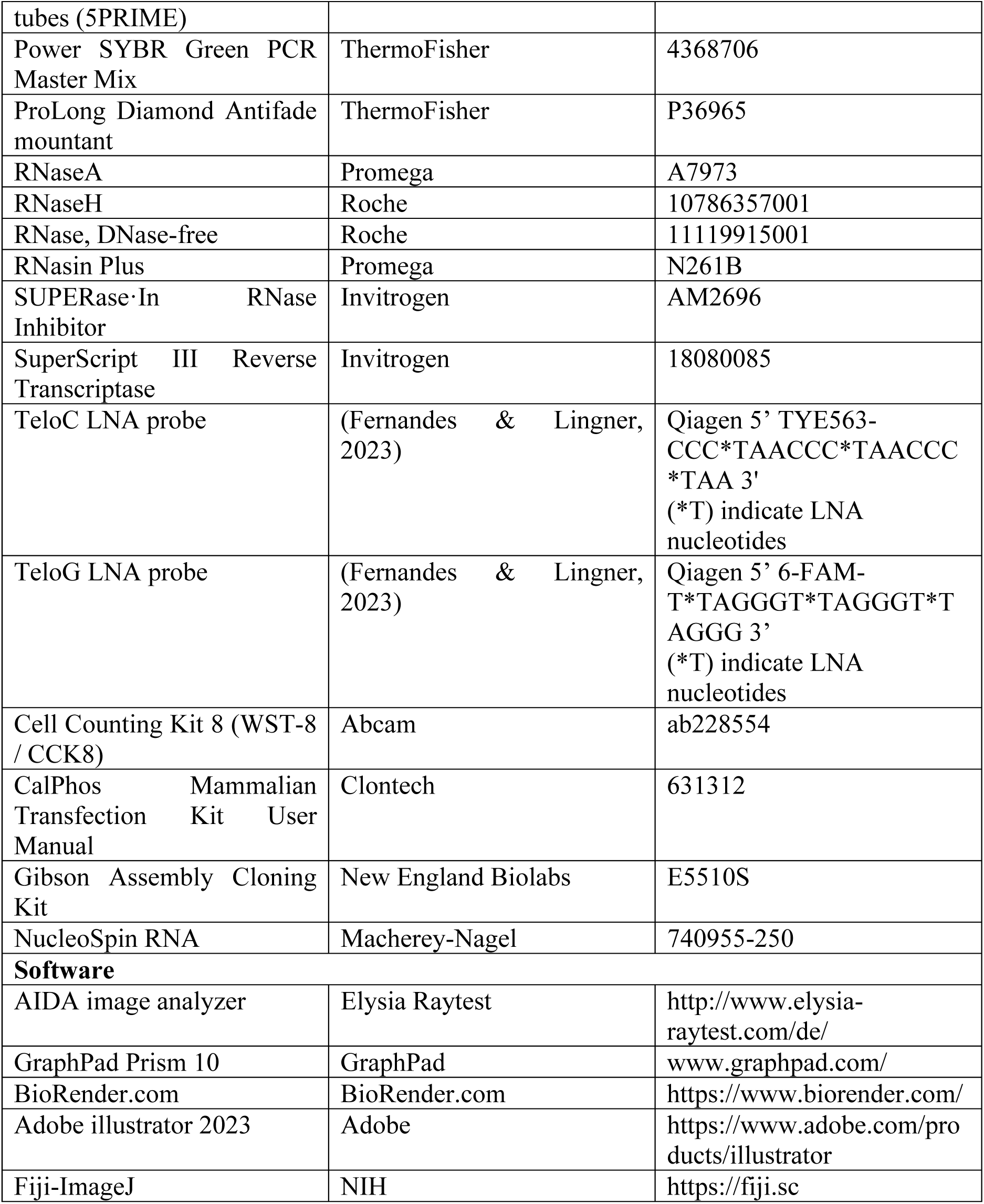

### Cell culture

HeLa and HEK293T cells were cultured in Dulbecco’s modified Eagle’s medium (DMEM) (Gibco), supplemented with 10% fetal bovine serum (FBS) and 100 U/mL penicillin/streptomycin. Cells were maintained at 37°C with 5% CO₂. The generation of HeLa cells with an average telomere length of 30 kb has been previously described (Cristofari & Lingner, 2006). The generation of TERRA-overexpressing cell lines is outlined below. Selection of dCas-VP64-expressing cells was carried out using 20 μg/mL blasticidin, gRNA-expressing cells were selected with 10 μg/mL puromycin, and cells expressing super-telomerase were selected with 250 μg/mL hygromycin B.

### Generation of TERRA overexpressing cell lines

To induce endogenous TERRA expression, the lentiviral vector Lenti-dCas-VP64_Blast and gRNA-containing plasmids (see Reagents and tools table) were introduced into HeLa cells. The gRNA-expressing plasmids contained a puromycin resistance gene. gRNAs were inserted into the modified vector via Gibson assembly (New England Biolabs), as described previously (Mali *et al*, 2013). The sequences of the gRNAs can be found in the Reagents and tools table.

### Lentivirus production and cell transduction

Lentiviral vectors were generated as previously described (Wiznerowicz *et al*, 2007). HEK293T cells (11 million) were seeded into 15 cm dishes a day prior to transfection. A combination of 55 μg pMD2-G, 102 μg PCMVR8.74 plasmids, and 157 μg transfer vector was transfected using the calcium phosphate method (CalPhos Mammalian Transfection Kit, Clontech). After overnight incubation, the medium was replaced, and the lentivirus-containing supernatant was collected over 24 hours in two intervals. The supernatant was filtered (0.22 μm) and concentrated by ultracentrifugation at 50,000 × g for 2 hours at 16°C. The viral pellet was resuspended in 500 μL PBS and stored at -80°C. Lentivirus titration was performed in HCT116 cells using qPCR, as previously described (Barde *et al*, 2010).

### siRNA transfection

siRNAs were used at a final concentration of 10 nM for the calcium phosphate transfection method. The day before transfection, 200,000 cells were seeded in 6-well plates, ensuring 30–40% confluency the following day. A 100 μL transfection mix was prepared, consisting of 125 mM CaCl₂, 110 nM siRNA, and 1× HBSS (pH 7.4) containing 50 mM HEPES, 280 mM NaCl, 1.19 mM Na₂HPO₄•2H₂O, and 10 mM KCl. The mix was incubated at RT for 1 min before being added to the cells in 1 mL of antibiotic-free DMEM supplemented with 10% FBS. Cells were harvested 72 hours post-transfection.

### Plasmid transfection

Plasmid transfection was carried out using Lipofectamine 2000 (Invitrogen) according to the manufacturer’s protocol with 4 μg of plasmid DNA. The day before transfection, 500,000 cells were seeded in 6-well plates to achieve 70-80% confluency by the time of transfection. A Lipofectamine pre-mix was prepared by combining 10 μL of Lipofectamine 2000 with 240 μL of Opti-MEM (Thermo Fisher) and incubating the mixture at room temperature for 5 minutes. Simultaneously, a DNA mix was prepared by mixing 4 μg of plasmid DNA with 250 μL of Opti-MEM. The Lipofectamine pre-mix was then added to the DNA mix and incubated at room temperature for 10 minutes. Finally, 500 μL of the transfection mix was added to the cells. Cells were harvested 48 hours after transfection.

### Western blotting

Cells were harvested by trypsinization and lysed in 2x Laemmli buffer. After heating at 95°C for 5 min, proteins were separated by SDS-PAGE and transferred to nitrocellulose membranes. Membranes were blocked with 3% BSA in PBS-Tween and incubated with primary antibodies overnight at 4°C. After washing, membranes were incubated with HRP-conjugated secondary antibodies, and signals were developed using chemiluminescence (ChemiGlow, BioTechne). Images were captured using a Fusion FX imaging system (Vilber).

### RNA purification, RNA dot blot and RT-qPCR

RNA was extracted from 3 to 5 million cells using the NucleoSpin RNA kit (Macherey-Nagel). The process included two on-column and one in-solution rDNase digestion (Macherey-Nagel). For RNA dot blot analysis, the purified RNA was treated with DNase-free RNase (Roche) as a control.

For RNA dot blot, samples were denatured at 65°C for 3 minutes and blotted onto a Hybond-XL membrane (Amersham) using a dot-blot apparatus (Bio-Rad). The RNA was UV-crosslinked to the membrane, which was then blocked in Church buffer (0.5 M NaHPO₄, 1 mM EDTA, pH 8.0, 1% BSA, 7% SDS) for at least 1 hour at 50°C. The membrane was hybridized overnight at 50°C with a ³²P-radiolabeled telomeric probe in Church buffer. After hybridization, the membrane was washed twice with 2x SSC, 0.5% SDS, and twice with 1x SSC, 0.5% SDS, for 15 minutes per wash. The radioactive signal was detected using a Typhoon Biomolecular Imager (GE) after exposure to a phosphorimager screen. Following signal detection, the membrane was stripped by incubation in boiling 0.1x SSC with 1% SDS at 55°C, repeated three times for 30 minutes each, and re-blocked in Church buffer for at least 1 hour at 55°C. A ³²P-radiolabeled 18S rRNA oligonucleotide probe was then hybridized overnight at 55°C, and the membrane was processed similarly as described for the telomeric probe.

For RT-qPCR analysis of TERRA levels, 3 μg of RNA was reverse transcribed using 200 U of SuperScript III reverse transcriptase (Thermo Fisher Scientific), along with GAPDH and TERRA reverse primers (see Reagents and tools table). The reverse transcription was performed at 55°C for 1 hour, followed by heat inactivation at 70°C for 15 minutes. Reactions without reverse transcriptase were included as controls. Five percent of the reaction was mixed with 2x Power SYBR Green PCR Master Mix (Thermo Fisher Scientific) and 0.5 μM forward and reverse qPCR primers (see Reagents and tools table). qPCR was performed in a QuantStudio 6 Flex Real-Time PCR system (Thermo Fisher Scientific) with the following cycling conditions: 95°C for 10 minutes, followed by 40 cycles of 95°C for 15 seconds, and 60°C for 1 minute for annealing and extension.

### Chromatin immunoprecipitation (ChIP)

10 million cells were harvested and crosslinked with 1% formaldehyde at RT for 10 min. Crosslinking was quenched by adding 1 M Tris-HCl (final concentration 250 mM) and rotating the mixture for 5 minutes at room temperature. The cells were then washed three times with cold 1X PBS. The cell pellets were flash-frozen in liquid nitrogen and stored at -80°C for further use. Frozen cell pellets were thawed on ice, resuspended in lysis buffer (1% SDS, 10 mM EDTA (pH 8.0), 50 mM Tris-HCl (pH 8.0), supplemented with cOmplete EDTA-free protease inhibitor (Roche)) and incubated on ice for 10 min. Samples were transferred to sonication vials with AFA fiber (Covaris), and sonicated with a Focused-Ultrasonicator (E220, Covaris) (10% duty factor, 140 W power, 200 cycles per burst, for 20 min), to shear DNA fragments of < 500 bp. Sonicated samples were centrifuged at 21,000 g for 10 min. Supernatants were collected and diluted with 1:10 in IP buffer (1.1% Triton, 1.2 mM EDTA pH 8.0, 16.7 mM Tris-HCl pH 8.0, 300 mM NaCl, supplemented with 1 tablet of protease inhibitor). Antibodies were added to each sample and incubated overnight at 4°C on a rotating wheel. Pre-blocked Sepharose G beads (incubated with ytRNA and IP buffer for 1 hour) were added to each sample, followed by incubation at 4°C for 4 hours. After incubation, the beads were pelleted by centrifugation at 400g for 2 min at 4°C, and the supernatant was discarded. The samples were washed by rotating on a wheel at 4°C for 5 min and centrifugation at 400 g for 2 min at 4°C, sequentially with low-salt (0.1% SDS, 1% Triton, 2 mM EDTA pH 8.0, 20 mM Tris pH 8.0, 300 mM NaCl), high-salt(0.1% SDS, 1% Triton, 2 mM EDTA pH 8.0, 20 mM Tris pH 8.0, 500 mM NaCl), LiCl (250 mM LiCl, 1% NP-40, 1% Na-deoxycholate, 1 mM EDTA pH 8.0, 10 mM Tris pH 8.0), and TE (1 mM EDTA pH 8.0, 10 mM Tris pH 8.0) buffers. Washed beads and input samples were resuspended in crosslink reversal buffer (20 mM Tris-HCl pH 8.0, 0.1% SDS, 0.1 M NaHCO₃, 0.5 mM EDTA) supplemented with 10 μg/mL RNase (DNase-free (Roche)) and incubated at 65°C overnight. DNA was purified using the NucleoSpin Gel and PCR Clean-up kit with NTB buffer (Macherey-Nagel) and eluted in 100 μL H2O. Samples were then analyzed by dot blot and qPCR (see below).

### DNA:RNA hybrid immunoprecipitation (DRIP)

DRIP was carried out as described previously (Glousker *et al*, 2022). Briefly, cells were harvested 72 hours after siRNA transfection or 48 hours after plasmid transfection via trypsinization. 1 × 10⁷ cells per condition were resuspended in 175 μL of ice-cold RLN buffer (50 mM Tris-HCl, pH 8.0, 140 mM NaCl, 1.5 mM MgCl₂, 0.5% NP-40, 1 mM dithiothreitol (DTT), and 100 U/mL RNasin Plus (Promega)) and incubated on ice for 5 minutes. Nuclei were pelleted by centrifugation at 300 g for 2 minutes at 4°C. The nuclear pellet was resuspended in 500 μL of RA1 buffer (NucleoSpin RNA purification kit, Macherey-Nagel) containing 1% 2-mercaptoethanol, then homogenized using a syringe with a 0.9 × 40 mm needle. The lysate was loaded into Phase Lock Gel Heavy tubes (5PRIME), mixed with 250 μL H₂O and 750 μL phenol-chloroform-isoamyl alcohol (25:24:1, pH 7.8-8.2), and centrifuged at 13,000 g for 5 minutes at room temperature. The aqueous phase was transferred to a new tube, and 750 μL of cold isopropanol and NaCl (final concentration 50 mM) were added. After mixing, the samples were incubated on ice for 30 minutes to precipitate nucleic acids, followed by centrifugation at 10,000 g for 30 minutes at 4°C. The resulting nucleic acid pellet was washed twice with 70% cold ethanol, air-dried, and dissolved in 130 μL of H₂O. The nucleic acids were then sonicated using a Covaris Focused-Ultrasonicator (E220, 10% duty factor, 140 W, 200 cycles per burst, 180 seconds) to generate fragments of 100–300 bp. The concentration of fragmented nucleic acids was measured using a NanoDrop spectrophotometer (ThermoFisher). For control, 10 μL of nucleic acids were digested with 10 μL of RNase H (1 U/μL, Roche) or H₂O in 150 μL of RNaseH buffer (20 mM HEPES-KOH pH 7.5, 50 mM NaCl, 10 mM MgCl₂, 1 mM DTT) and incubated at 37°C for 90 minutes. The reaction was stopped by adding 2 μL of 0.5 M EDTA (pH 8.0). Nucleic acids were diluted 1:10 in DIP-1 buffer (10 mM HEPES-KOH pH 7.5, 275 mM NaCl, 0.1% SDS, 0.1% Na-deoxycholate, 1% Triton X-100) and pre-cleared with 40 μL of Sepharose Protein G beads (Cytiva) for 1 hour at 4°C. DRIP was carried out using 30 μg of nucleic acids from HeLa cells with 3 μg of the S9.6 antibody (Kerafast) or mouse IgG. After incubation overnight at 4°C on a rotating wheel, samples were washed with buffer DIP-2 (50 mM HEPES–KOH pH 7.5, 140 mM NaCl, 1 mM EDTA pH 8.0, 1% Triton X-100, 0.1% Na-deoxycholate), buffer DIP-3 (50 mM HEPES–KOH pH 7.5, 500 mM NaCl, 1 mM EDTA pH 8.0, 1% Triton-X100, 0.1% Na-deoxycholate), buffer DIP-4 (10 mM Tris–HCl pH 8.0, 1 mM EDTA pH 8.0, 250 mM LiCl, 1% NP-40, 1% Na-deoxycholate), and TE buffer (10 mM Tris–HCl pH 8.0, 1 mM EDTA pH 8.0). Immunoprecipitated samples and inputs were eluted in 100 μL of crosslink reversal buffer (20 mM Tris-HCl pH 8.0, 0.1% SDS, 0.1 M NaHCO₃, 0.5 mM EDTA) containing 10 μg/mL RNase (DNase-free (Roche)), and incubated at 65°C overnight. DNA was purified using the QIAquick PCR Purification Kit (Qiagen) and eluted in 100 μL of H₂O. The purified samples were then analyzed by dot blot or qPCR.

### qPCR analysis of ChIP and DRIP samples

Each qPCR reaction consisted of 1 μL of purified DNA (from immunoprecipitated or diluted input samples as described), 5 μL of Power SYBR Green PCR Master Mix (Thermo Fisher Scientific), 1 μM of forward and reverse primers (see Reagents and tools table), and H₂O to a final volume of 10 μL. Each sample was run in technical duplicates. The qPCR was performed with an initial denaturation at 95°C for 10 minutes, followed by 40 cycles of 95°C for 15 seconds and annealing/extension at 60°C for 1 minute, using a QuantStudio 6 Flex Real-Time PCR system (Thermo Fisher Scientific). Serial dilutions (with factors of 5 and 50) of each input sample were included to generate a standard curve via regression analysis. Based on the input equation, the corresponding immunoprecipitation values were calculated as a percentage of input.

### Dot blot analysis of ChIP and DRIP samples

Input samples (from ChIP and DRIP experiments) were appropriately diluted to ensure the DNA concentration in each immunoprecipitated sample remained within the assay’s linear dynamic range. Both purified DNA from diluted input and immunoprecipitated samples were heated at 95°C for 5 minutes, then immediately cooled on ice. The samples were then blotted onto a Hybond-XL membrane (Amersham) using a dot blot apparatus (Bio-Rad) and UV-crosslinked to the membrane. The membrane was denatured in a solution of 0.5 M NaOH and 1.5 M NaCl for 15 minutes at room temperature on a shaker, followed by neutralization in 0.5 M Tris-Cl (pH 7.0) and 1.5 M NaCl for 10 minutes. The membrane was blocked in Church buffer (0.5 M NaHPO₄, 1 mM EDTA, pH 8.0, 1% BSA, 7% SDS) at 65°C for at least 1 hour. Hybridization was performed overnight at 65°C with a ³²P-radiolabeled TeloC probe in Church buffer. The membrane was then washed three times for 30 minutes at 65°C in 1× SSC containing 0.5% SDS, and exposed to a phosphorimager screen. The radioactive signal was detected using a Typhoon Biomolecular Imager (GE).

### Immunofluorescence and telomeric fluorescence in situ hybridization (IF-FISH)

For zeocin treatment, cells were incubated with 100 µg/mL of zeocin for 3 hours. Cells were cultured on round coverslips, washed twice with PBS, and fixed in 4% paraformaldehyde for 10 minutes at room temperature. After fixation, cells were permeabilized with a detergent solution (0.1% Triton X-100, 0.02% SDS in PBS) and blocked with 2% BSA in PBS for 10 minutes. Primary antibodies were applied in a blocking solution (10% normal goat serum, 2% BSA in PBS) and incubated overnight at 4°C. Secondary antibodies were applied for 30 minutes at room temperature after multiple washes. Coverslips were then fixed again in 4% paraformaldehyde, washed, and dehydrated with a graded ethanol series (70%, 95%, and 100%). For FISH, hybridization was performed using 100 nM Cy3-[CCCTAA]3 PNA probe in a hybridization mix (10 mM Tris, 70% formamide, 0.5% blocking reagent) at 80°C for 3 minutes, followed by incubation at room temperature for 3 hours. Post-hybridization washes were performed twice with wash buffer 1 (10 mM Tris, 70% formamide) and three times with wash buffer 2 (0.1 M Tris, 0.15 M NaCl, 0.08% Tween-20). DAPI was added to the second wash step of wash buffer 2. Coverslips were mounted with Vectashield, and images were captured with a Leica SP8 confocal microscope equipped with a 63 ×/ 1.40 oil objective and a DFC 7000 GT camera.

### Telomeric FISH on metaphase chromosomes

Telomeric FISH was performed as previously described (Fernandes & Lingner, 2023). Cells were treated with 0.05 μg/mL demecolcine for 2 hours before being harvested and subjected to hypotonic treatment (0.056 M KCl) at 37°C for 7 minutes. After fixation in cold methanol acetic acid (3:1) overnight, cells were spread onto slides and processed for FISH staining, as described above, using a 100 nM Cy3-[CCCTAA]3 PNA probe. Images were captured using a Zeiss Axioplan or Leica SP8 confocal microscope with a Upright Zeiss Axioplan equipped with a 100x/1.40 oil objective.

### Chromosome Orientation (CO)-FISH on metaphase chromosomes

CO-FISH was carried out as previously described (Fernandes & Lingner, 2023). Cells were incubated with BrdU/BrdC (3:1, final concentration of 10 μM) for 15 hours and treated with 0.1 μg/mL demecolcine for 2 hours. After hypotonic treatment, cells were fixed and spread onto glass slides. After rehydration, cells were treated with RNaseA (Promega), stained with Hoechst 33258, and exposed to UV light. Following exonuclease digestion and paraformaldehyde fixation, hybridization was performed using TYE563- TeloC LNA and 6-FAM-TeloG LNA probes (Qiagen). Post-hybridization washes were performed, and slides were mounted with ProLong Diamond Antifade mountant. Images were acquired using a Leica SP8 confocal microscope equipped with a 63 ×/ 1.40 oil objective and a DFC 7000 GT camera.

### WST-8 cell proliferation assay

The WST-8 cell proliferation assay was conducted according to the manufacturer’s protocol (Abcam, Ab228554). Briefly, 15,000 cells were seeded in 24-well plates. At 24, 48, 72, and 96 hours (corresponding to days 2, 3, 4, and 5), the culture media were replaced with 500 µL of WST-8 solution (450 µL of pre-warmed fresh media mixed with 50 µL of WST-8 reagent). After incubating the plates at 37°C for 1 hour, 200 µL of the supernatant was transferred to a 96-well plate (Falcon, 351172), and the absorbance was measured at 460 nm using a plate reader. siRNA transfection was performed on day 2 using the calcium phosphate transfection method.

### Software

Image analysis was done using ImageJ, dot blots were analyzed with Aida Image Analyzer, and statistical analyses and graph preparation were performed using GraphPad Prism. Illustrations were created using Adobe Illustrator 2023 and BioRender.com.

## Acknowledgements

We thank Eftychia Kyriacou, Satyajeet Rao, and Galina Glousker for comments on the manuscript. This work was supported by the Swiss National Science Foundation (SNSF) [310030_214833] and the SNSF-funded National Centre of Competence in Research RNA and Disease Network [205601]. PRN was recipient of PhD fellowship from the Boehringer Ingelheim Fonds.

## Author contributions

**Suna In**: Conceptualization; Formal analysis; Data curation; Validation; Investigation; Visualization; Methodology; Writing—original draft; Writing—review and editing. **Patricia Renck Nunes**: Methodology; Writing—review and editing. **Rita Valador Fernandes**: Methodology; **Joachim Lingner**: Conceptualization; Data curation; Supervision; Funding acquisition; Investigation; Writing—original draft; Project administration; Writing—review and editing.

## Supplementary Material

**Figure S1.**
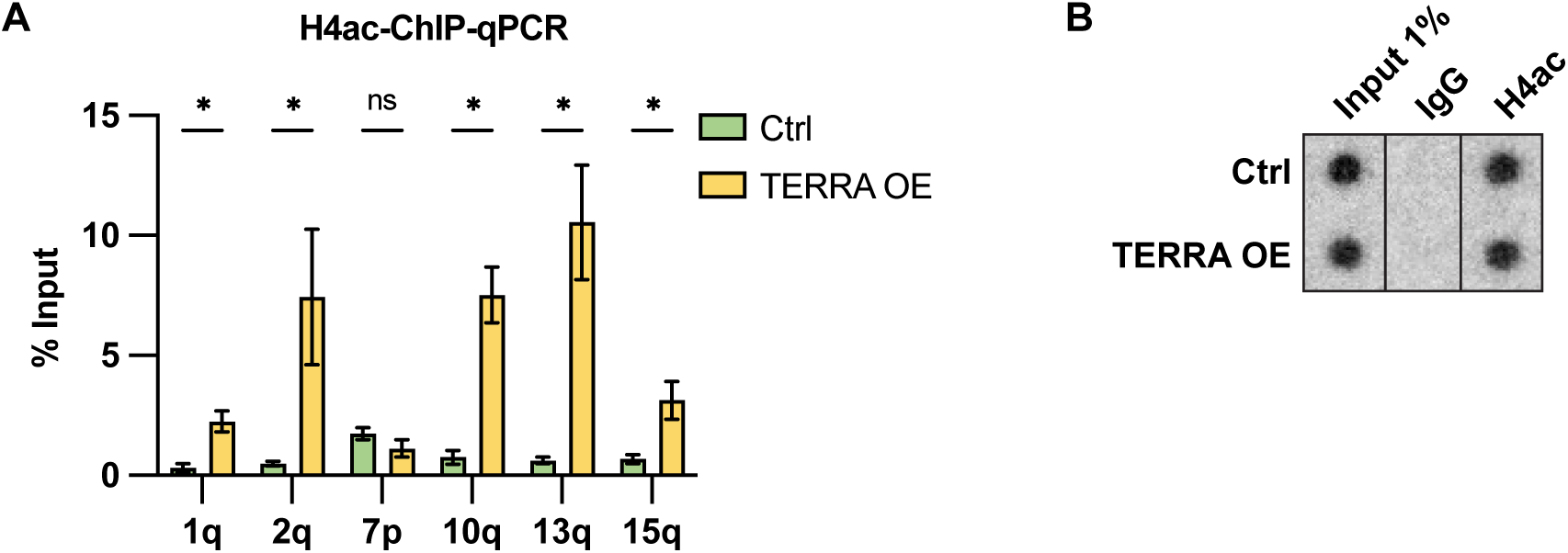
Induction of endogenous TERRA by modified CRISPR-Cas9 system, related to Figure 1. (A-B) ChIP assay using H4 acetylation antibody. ChIP samples and inputs were treated with RNase (DNase-free) and analyzed by qPCR with indicated subtelomeric primers (A) or by DNA dot blot with a ^32^P-radiolabeled telomeric probe (B).

**Figure S2.**
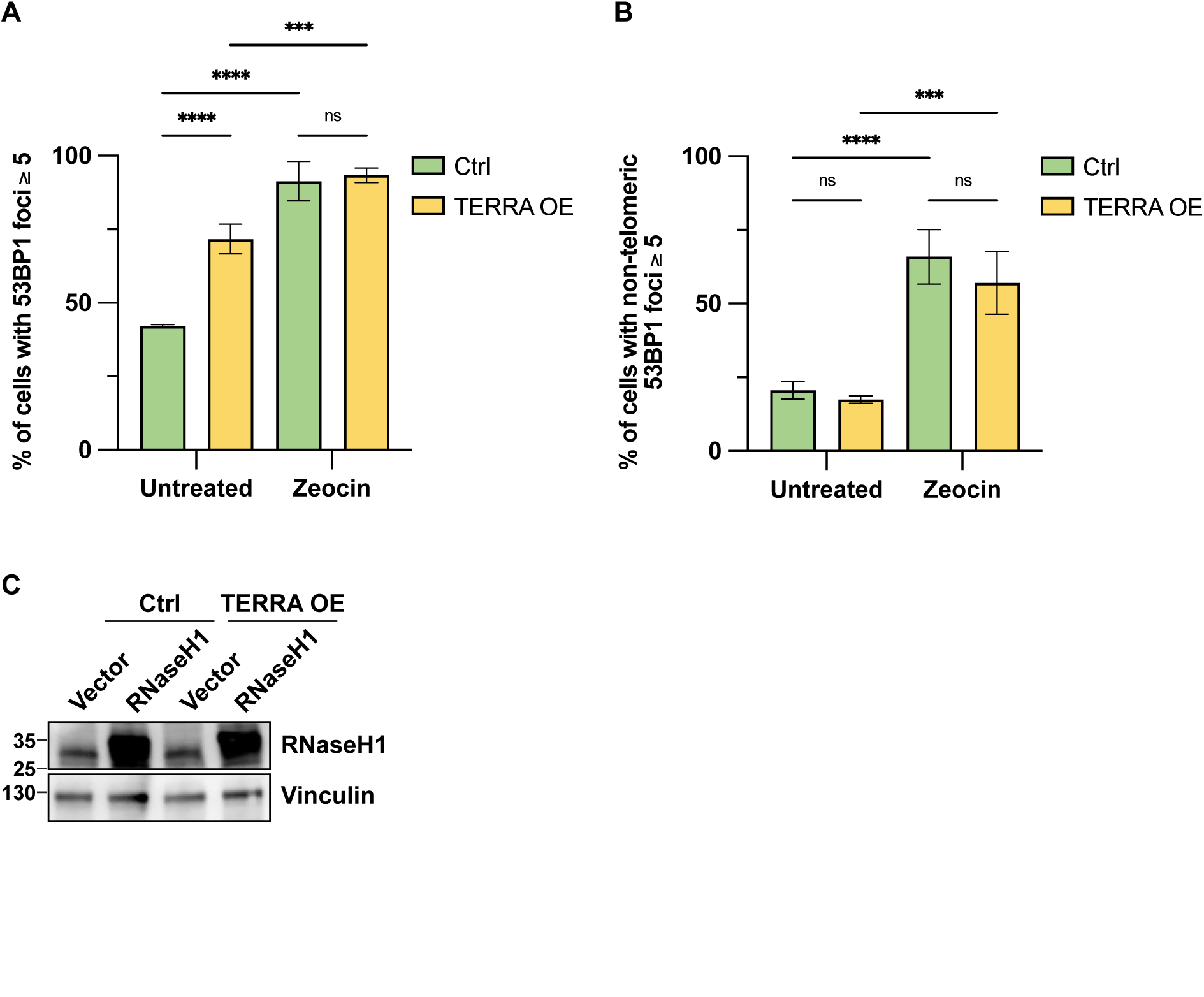
DNA damage upon TERRA overexpression and zeocin treatment, related to Figure 3. Quantification of the number of cells with ≥ 5 53BP1 foci. Data represent mean ± s.d., from three independent biological replicates. Two-way analysis of variance (ANOVA) with uncorrected Fisher’s least significant difference (LSD) test was applied: **** P ≤0.0001, *** P ≤ 0.001, ns indicates non-significance (P > 0.05). (A) Quantification of the number of cells with ≥ 5 53BP1 foci that are not colocalizing with telomeres. Data represent mean ± s.d., from three independent biological replicates. Two-way analysis of variance (ANOVA) with uncorrected Fisher’s least significant difference (LSD) test was applied: **** P ≤0.0001, *** P ≤ 0.001, ns indicates non-significance (P > 0.05). (B) Western blot analysis upon ectopic expression of RNaseH1 in control and TERRA overexpressing HeLa cells.

**Figure S3.**
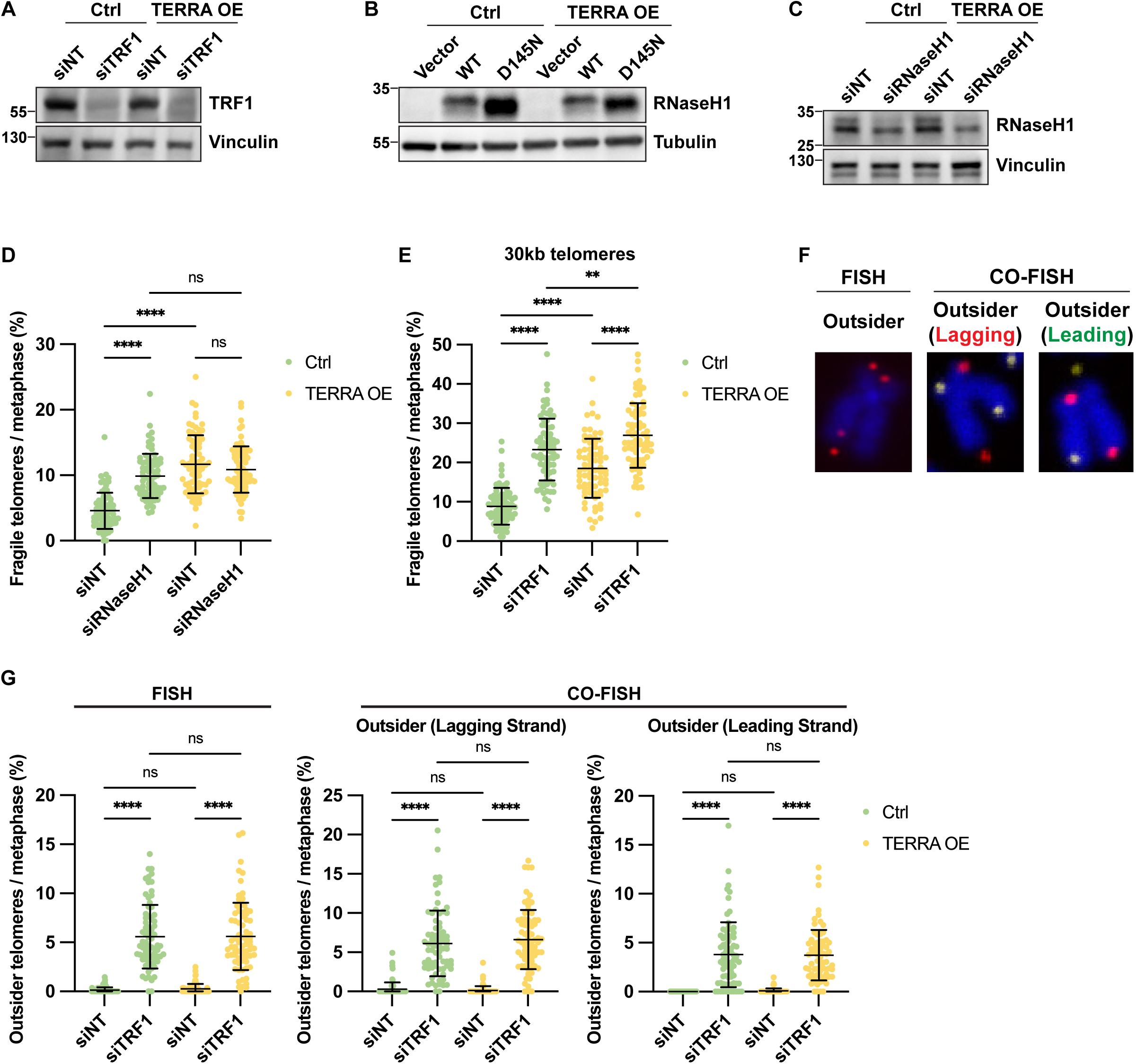
Fragile and outsider telomeres, related to Figure 4. (A) Western blot analysis upon depletion of TRF1 in control and TERRA overexpressing HeLa cells. (B) Western blot analysis upon ectopic expression of RNaseH1 WT or D145N mutant in control and TERRA overexpressing HeLa cells. (C) Western blot analysis upon depletion of RNaseH1 in control and TERRA overexpressing HeLa cells. (D) Quantification of telomere fragility upon depletion of RNaseH1 in control and TERRA overexpressing HeLa cells. At least 25 metaphases were analyzed per condition per replicate, and three independent biological replicates were performed. Horizontal lines and error bars represent mean ± s.d. Two-way analysis of variance (ANOVA) with Tukey’s multiple comparisons test was applied: **** P ≤0.0001, ns indicates non-significance (P > 0.05). (E) Representative images of outsider telomeres by FISH (left) or CO-FISH (middle and right). Metaphase spreads were stained with telomeric (CCCTAA)_3_-FISH probe (red) and DAPI (blue) (left), with TYE563-TeloC LNA probe (red), FAM-TeloG LNA probe (yellow), and DAPI (blue) (middle and right). (F) Quantification of telomere fragility upon depletion of TRF1 in control and TERRA overexpressing HeLa cells with 30 kb telomeres. At least 25 metaphases were analyzed per condition per replicate, and three independent biological replicates were performed. Horizontal lines and error bars represent mean ± s.d. Two-way analysis of variance (ANOVA) with Tukey’s multiple comparisons test was applied: **** P ≤0.0001, ** P ≤ 0.01. (G) Quantification of outsider telomeres upon depletion of TRF1 in control and TERRA overexpressing HeLa cells with 30 kb telomeres. Metaphases from the same samples were analyzed by FISH (left) and CO-FISH (middle and right). At least 25 metaphases were analyzed per condition per replicate, and three independent biological replicates were performed. Horizontal lines and error bars represent mean ± s.d. Two-way analysis of variance (ANOVA) with Tukey’s multiple comparisons test was applied: **** P ≤0.0001, ns indicates non-significance (P > 0.05).

**Figure S4.**
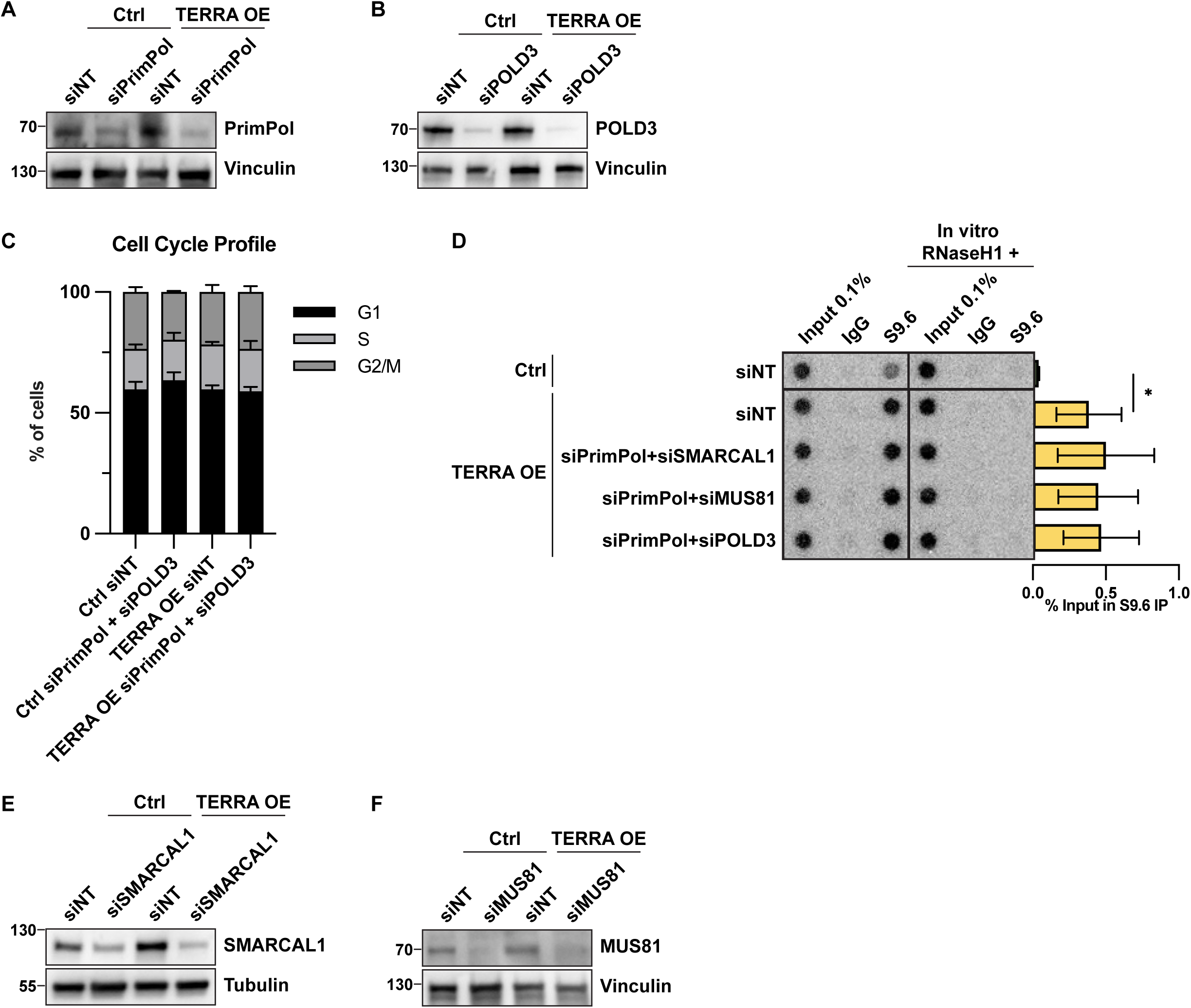
Western blot analyses of depletion of proteins involved in telomere fragility, related to Figure 5. (A) Western blot analysis upon depletion of PrimPol in control and TERRA overexpressing HeLa cells. (B) Western blot analysis upon depletion of POLD3 in control and TERRA overexpressing HeLa cells. (C) Cell cycle profiles upon PrimPol and POLD3 depletion. DNA content was analyzed by flow cytometry analysis of fixed DAPI-stained cells. (D) DRIP assay using S9.6 antibody. DRIP samples and inputs were treated with RNase (DNase-free) and analyzed by DNA dot blot with a ^32^P-radiolabeled telomeric probe. As a negative control, samples were treated in vitro with RNaseH1 prior to immunoprecipitation and analyzed in parallel. Data represent mean ± s.d., from five independent biological replicates. Unpaired t-test was applied: * P ≤ 0.05 (D) Data represent mean ± s.d., from three independent biological replicates. Multiple unpaired t-test was applied: * P ≤ 0.05. (E) Western blot analysis upon depletion of SMARCAL1 in control and TERRA overexpressing HeLa cells. (F) Western blot analysis upon depletion of MUS81 in control and TERRA overexpressing HeLa cells.

**Figure S5.**
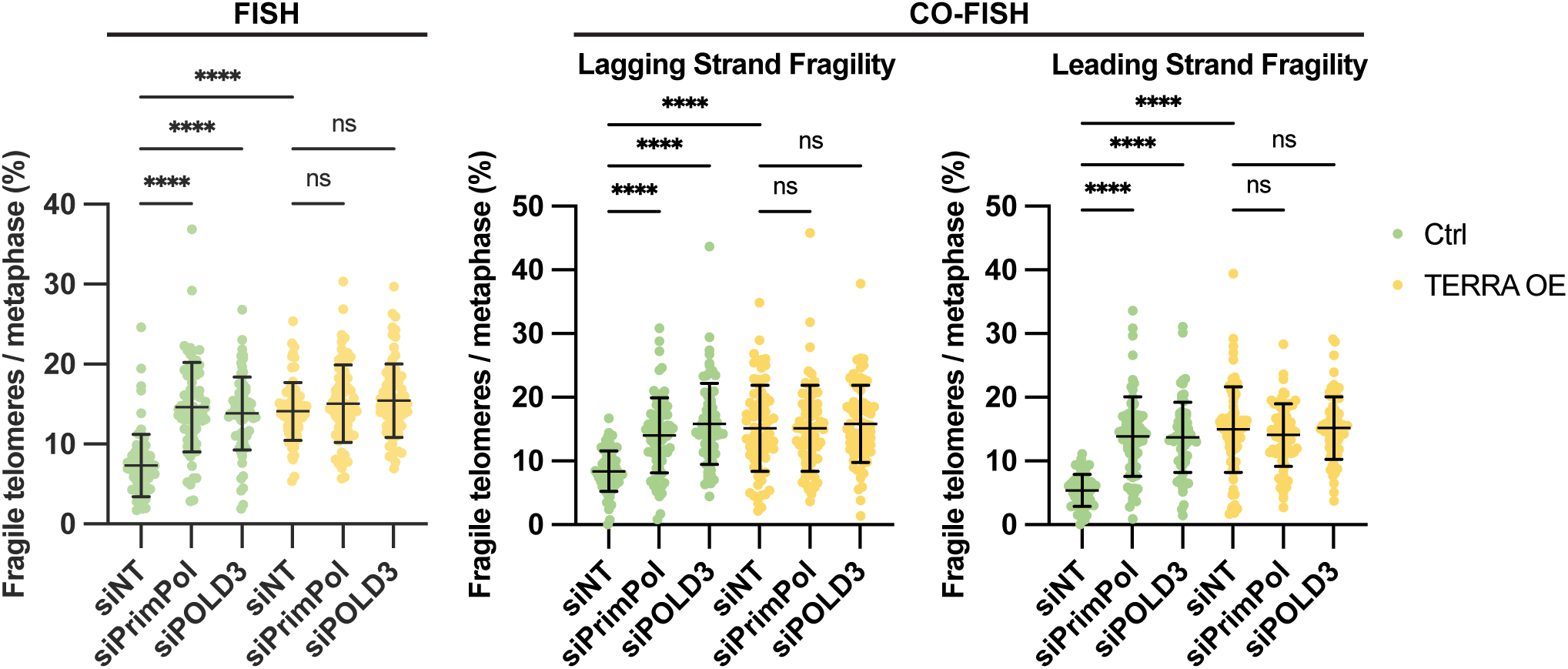
Impacts of depletion of PrimPol, POLD3, SMARCAL1, and MUS81, related to Figure 6. Quantification of telomere fragility upon depletion of PrimPol or POLD3 in control and TERRA overexpressing HeLa cells with 30 kb telomeres. Metaphases from the same samples were analyzed by FISH (left) and CO-FISH (middle and right). At least 25 metaphases were analyzed per condition per replicate, and three independent biological replicates were performed. Horizontal lines and error bars represent mean ± s.d. Two-way analysis of variance (ANOVA) with Tukey’s multiple comparisons test was applied: **** P ≤0.0001, ns indicates non-significance (P > 0.05).

